# PDR6-mediated camalexin efflux and disease resistance are regulated through direct phosphorylation by the kinases OXI1 and AGC2-2

**DOI:** 10.1101/2022.02.08.479400

**Authors:** Guopeng Miao, Juan Han, Chang-xin Liu, Jian Liu, Cheng-run Wang, Shun-chang Wang

**Author notes:** **Correspondence:** Corresponding Author: Guopeng Miao.

## Abstract

Plant immune signaling largely relies on post-translational modifications to establish a rapid and appropriate defense response to different pathogen types and infection pressure. Specific pleiotropic drug resistance (PDR) transporters can transport secondary metabolites to contribute to pathogen invasion resistance. However, the establishment of the post-translational regulation of PDR transporters that efflux secondary metabolites is unclear. In this study, by detecting the camalexin contents on the leaf surfaces of mutants and overexpression lines, two AGC kinases, namely, OXI1 and its closest homologue AGC2-2, were found to be related to extracellular camalexin secretion. The overexpression of *OXI1* or *AGC2-2* resulted in an increase in camalexin contents on the leaf surface and a decrease in camalexin contents in the leaf interior. These effects increased the resistance of the transgenic lines to surface-inoculated *Pseudomonas syringae* and *Botrytis cinerea*. Through *in vitro* kinase assay and *in vivo* phosphorylation level detection, we confirmed that the two kinases were related to the phosphorylation modification of PDR6. Pull-down assays, bimolecular fluorescence complementation, and rapamycin-dependent delocalization assays indicated the existence of direct protein–protein interaction between the two kinases and PDR6. By using LC–MS/MS, we also identified the PDR6 phosphorylation sites that were modified by the two kinases *in vitro*. Through the expression of the dephosphorylated variants of PDR6 in the mutant background, the action site S31 of OXI1 and the action sites S33 and S827 of AGC2-2 were found to have positive effects on the efflux activity of PDR6. In addition, T832, the action site of OXI1, may contribute to the stability of PDR6 on the plasma membrane.

## Introduction

Plants resist pathogen attacks via two kinds of innate immune responses: one is initiated by the recognition of pathogen-associated molecular patterns or damage-associated molecular patterns *via* cell-surface-localized pattern recognition receptors and leads to pattern-triggered immunity (PTI), and the other is initiated by the recognition of effectors through intracellular nucleotide-binding domain leucine-rich repeat-containing receptors and leads to effector-triggered immunity[1]. Following the recognition of patterns or effectors, such as chitin, plants deploy a variety of mechanisms to prevent or limit microbial infection; these mechanisms involve changes in ion fluxes, such as an increase in cytoplasmic calcium concentration, the production of reactive oxygen species (ROS), the activation of mitogen-activated protein kinase (MAPK) gene families, and alterations in plant phytohormone and metabolite status[2]. Among these mechanisms, chemicals synthesized on the plant surface can inhibit the colonization and penetration of plant tissues or cells by potential pathogens. These compounds include cell-wall-degrading enzymes (e.g., chitinase and glucanase); antimicrobial proteins (e.g., thionine and defensin);[3] and low-molecular-weight secondary metabolites, which are secreted outside the cells *via* transporters that not only defend against pathogens[4] but also limit the cytotoxic effects of phytoalexin overaccumulation in plant cells[5].

At present, two types of transporters are mainly involved in the transport of plant secondary metabolites: ATP-binding cassette (ABC)[6] and multidrug and toxic compound extrusion (MATE) transporters[7]. As a subclass of ABC transporters, the pleiotropic drug resistance (PDR)/ATP binding cassette G (ABCG) transporters, which have a full molecular structure that is composed of two nucleotide-binding domains (NBDs) and two transmembrane domains (TMDs) and are arranged in the orientation of NBD–TMD–NBD–TMD, are involved in important processes that influence plant fitness; these processes include pathogen response, diffusion barrier formation, or phytohormone transport[8]. Yusuke et al.[9] found that tobacco transports capsidiol outside the cell through two PDR transporters, namely, NbABCG1 and NbABCG2, to resist the invasion of *Phytophthora infestans* into the leaves. In *Nicotiana plumbaginifolia*, after fungal pathogen challenge, NpPDR1 secretes diterpenoids, which may possibly be sclareolide and sclareol, for host protection[10, 11]. Camalexin (3-thiazol-2′-yl-indole), the characteristic phytoalexin of *Arabidopsis thaliana*, is induced by a great variety of plant pathogens. The deletion mutant of the PDR6 transporter in *Arabidopsis* is highly sensitive to the diterpenoid sclareol and significantly reduces the amounts of camalexin secreted on the leaf surface[12]. However, He et al.[13] showed that camalexin efflux is mainly mediated by PDR8/PEN3 and PDR12 but not by PDR6.

Although many studies have reported that ABC transporters can transport secondary metabolites to the outside of the cell to resist pathogen invasion, the mechanism of transporter activity regulation to achieve this function in plants has been less reported than that in yeast and mammals[14]. The phosphorylation of plant transporters can change the transport activity, stability, and intracellular transport of proteins and plays an important regulatory role in the transport of many plant components, such as sugar[15], auxin[16], inorganic nutrients[17, 18], and defense-related compounds[19]. However, among all plant ABC transporters, the upstream kinase signal pathway has only been identified in limited cases. The two auxin transporters ABCB1 and ABCB19 can be phosphorylated by the AGC (orthologs of mammalian PKA, PKG, and PKC) kinase family members PID[20] and PHOT1[21], respectively. Further phosphorylation proteomic analysis has found that the amino acid residue S634 in ABCB1 is likely to be the phosphorylation site of PID with an important role in substrate binding and transport activity[20].

The *Arabidopsis* AGC kinase family has 39 members, which are divided into eight categories; PHOT1 and PID belong to the AGC4 and AGC3 subclasses, respectively[22]. The functionally identified AGC kinases, except for the AGC2 subclass kinase oxidative signal-inducible 1 (OXI1/AGC2-1), are mainly involved in light signaling and cell polarity regulation. OXI1 is necessary for oxidative burst-mediated signaling[23] and plant immunity against *Pseudomonas syringae* and *Hyaloperonospora parasitica*[24]. It also participates in MAPK cascade signal transduction[25]. As the closest homolog of OXI1[22], AGC2-2 has been shown to have redundant yet presumably different functions[26].

Plant immune signaling largely relies on post-translational modifications to establish a rapid and appropriate defense response to pathogen type and infection pressure[27, 28]. However, how post-translational regulation is established on the basis of PDR transporters that efflux secondary metabolites is unclear. In this study, the activity of PDR6 was proven to be regulated by OXI1 and AGC2-2 *via* direct phosphorylation modifications. Our results would help reveal how plants transmit signals to transporters to resist pathogen invasion quickly.

## Results

### OXI1 and AGC2-2 are involved in camalexin efflux

The genes coexpressed with the three transporters PDR6, PDR8, and PDR12 were obtained from the ATTED-II database[29] to identify the candidate AGC kinase(s) involved in the transportation of camalexin. After Gene Ontology analysis, OXI1 was screened out manually because it was coexpressed with PDR6 and PDR12 (data not shown) and selected as a candidate kinase along with its closest homolog AGC2-2. First, the levels of camalexin secretion in the wild-type *Arabidopsis* and the mutants were examined in detail. Camalexin accumulation on the leaf surfaces of *oxi1* and *agc2-2* was significantly lower than that on the leaf surfaces of Columbia-0 (Col-0). Camalexin secretion by the *oxi1* and *agc2-2* mutants was 58.7% and 63.6% lower than that by Col-0, respectively, after 48 h of chitin elicitation (**Fig. 1A–B**). A double mutant of *oxi1* and *agc2-2*, which may remain heterozygous for one insertion and homozygous for the other[30], showed even lower camalexin contents than that of single mutant (**Fig. 1A–B**). This result indicated that the two kinases might play overlapping yet independent roles in camalexin secretion.

**Fig.1.**
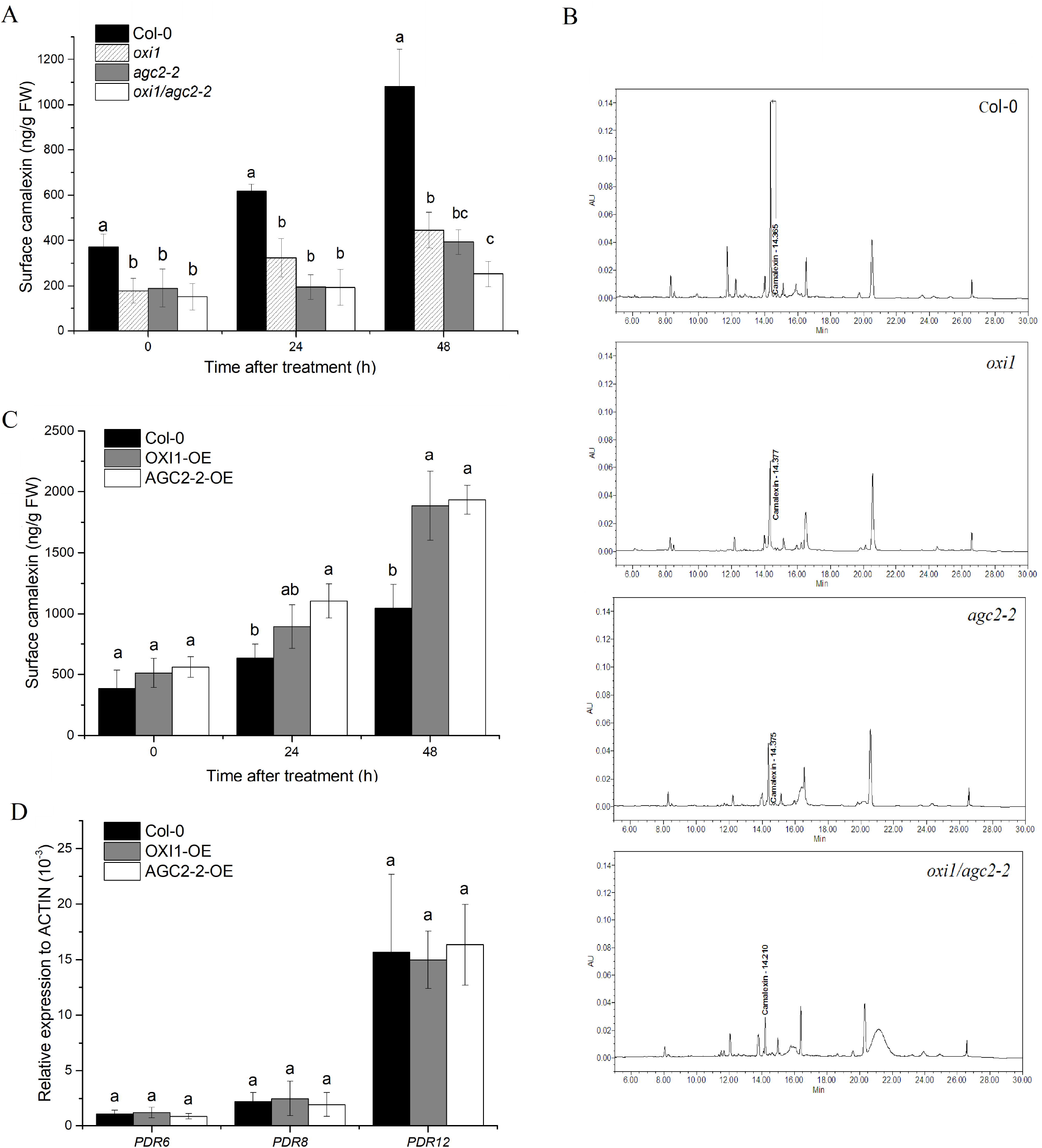
OXI1 and AGC2-2 are related to camalexin secretion on the leaf surface but not to the transcriptional regulation of transporters. Camalexin contents on the leaf surfaces of the mutants oxi1, agc2-2, and oxi1/agc2-2 were lower than those on the leaf surfaces of wild-type Col-0 before and after chitin treatment (**A**). Compounds in the leaf extrudates were separated through HPLC, and camalexin was identified by using an external standard and UV absorption spectra (**B**). After chitin induction, *OXI1* and *AGC2-2* overexpression increased camalexin secretion (**C**). The transcription of the three transporters PDR6, PDR8, and PDR12 was unaffected by the overexpression of *OXI1* or *AGC2-2* (**D**). Data are presented as mean ± SD, n = 3. Letters on scale bars indicate significant differences in accordance with Duncan’s multiple range test at p < 0.01. OE: overexpression.

Stable transgenic lines with *OXI1* or *AGC2-2* overexpression driven by the cauliflower mosaic virus 35S promoter were established to further confirm the relationship among OXI1, AGC2-2, and camalexin secretion. The overexpression of *OXI1* or *AGC2-2* did not influence plant growth (**Fig. S1**). Under unchallenged conditions, i.e., 0 h of treatment, no significant difference in camalexin secretion between the overexpression lines and Col-0 was observed. However, at 48 h post-treatment, camalexin secretion by the *OXI1* and *AGC2-2* overexpression lines increased to approximately 1.80- and 1.85-fold that by the control, respectively (**Fig. 1C**). In the transgenic lines, the transcript levels of the three transporters PDR6, PDR8, and PDR12 did not show significant changes (**Fig. 1D**). This result illustrated that OXI1 and AGC2-2 did not regulate the transcription of the transporter genes.

### OXI1 and AGC2-2 positively regulate resistance to the pathogens *P. syringae* and *Botrytis cinerea*

Camalexin is involved in defense against various pathogens[31]. Given that OXI1 and AGC2-2 are required for the full secretion of camalexin onto the leaf surfaces of *Arabidopsis*, their participation in disease resistance was further investigated. Previously, Petersen et al.[24] demonstrated that OXI1 is required for full resistance to *P. syringae*. However, the overexpression experiments showed conflicting results. We speculated that this discrepancy may be caused by the decrement in camalexin in cells due to the overexpression of *OXI1* but not the disruption of OXI1 function as suggested by the authors. *P. syringae* was inoculated through leaf dipping but not through infiltration to confirm our hypothesis. The results showed clear consistency between the reduced expression and overexpression of *OXI1* and *AGC2-2*: At 48 h after inoculation, susceptibility increased in the *oxi1, agc2-2*, and *oxi1/agc2-2* mutants but decreased in the overexpression lines (**Fig. 2A–B**). The decreased camalexin contents within the cells further confirmed our speculation (**Fig. 2C**). Notably, the total contents of camalexin (combined leaf surface and interior) in the overexpression lines were lower than those in Col-0 (**Fig. 1C** and **2C**).

**Fig.2.**
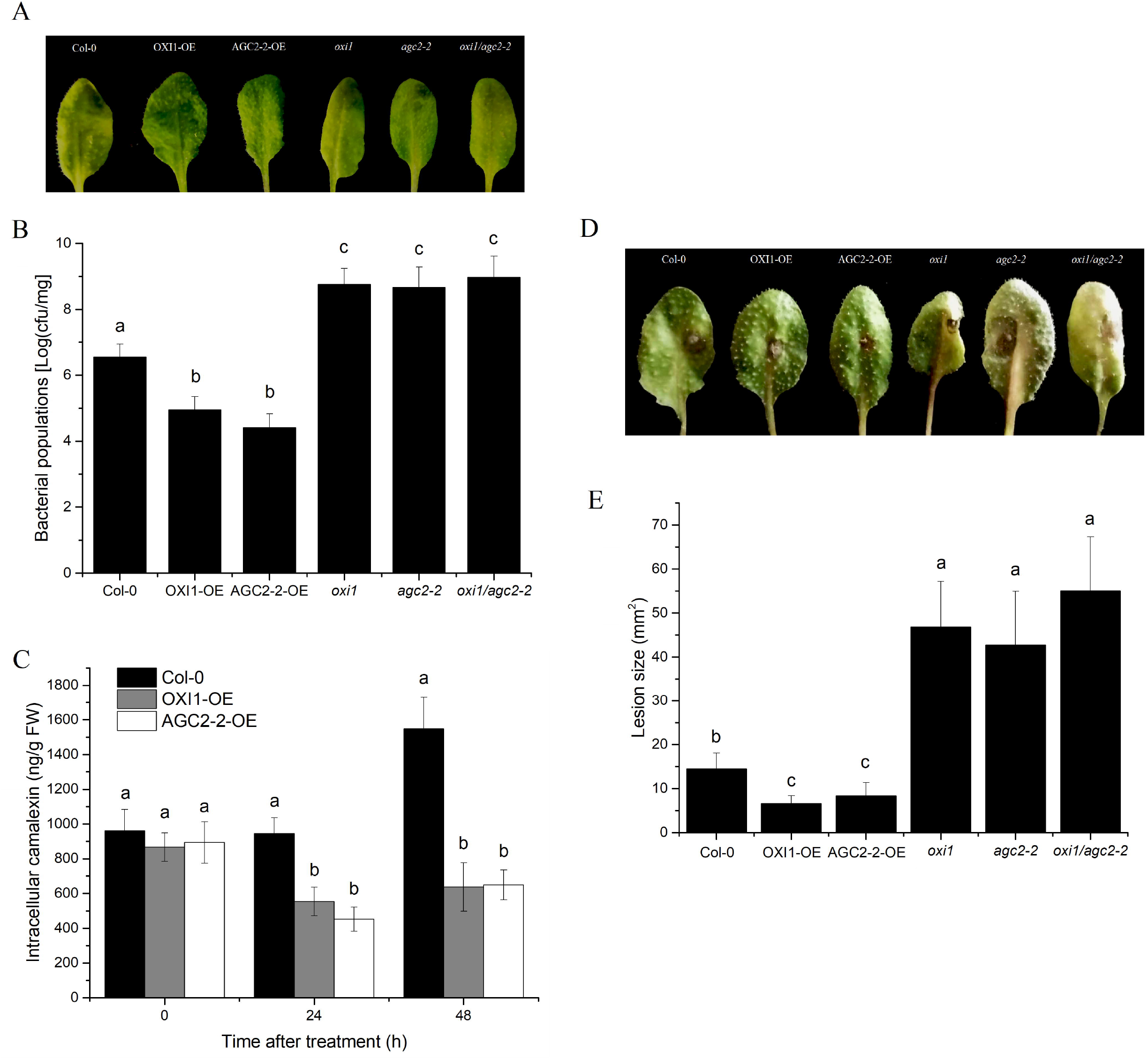
OXI1 and AGC2-2 positively regulate disease resistance to the pathogen *P. syringae* pv. *tomato* DC 3000 and *B. cinerea*. After 3 days of *P. syringae* infiltration in the leaves of the tested *Arabidopsis*, disease symptoms were recorded (**A**) and bacterial populations were counted (**B**). The intracellular contents of camalexin in the overexpression lines of OXI1 and AGC2-2 were lower than those in Col-0 after *P. syringae* infiltration (**C**). The disease resistance of genetically manipulated lines to *B. cinerea* was examined, and disease symptoms (**D**) and lesion sizes (**E**) were recorded at 3 days after inoculation. Data are presented as mean ± SD, n = 3–6. Letters on scale bars indicate significant differences in accordance with Duncan’s multiple range test at p < 0.01. OE: overexpression.

Considering that the impaired secretion of camalexin has been reported to be capable of increasing susceptibility to the necrotrophic pathogen *Botrytis cinerea*[12, 13], the disease resistance of the genetically manipulated lines to this pathogen was also examined. At 3 days after inoculation, resistance responses to *B. cinerea* that were similar to those to *P. syringea* were observed (**Fig. 2D–E**).

### Phosphorylation of PDR6 is related to OXI1 and AGC2-2

Previous studies have debated on which transporter mediates the extracellular secretion of camalexin[12, 13]. Therefore, we performed a validation experiment. Our results showed that the amounts of camalexin secreted by the *pdr6* and *pdr8* mutants were significantly lower by 48.4% and 27.5%, respectively, than those secreted by Col-0, whereas the PDR12 mutant did not show any effects (**Fig. S2**). Given that PDR6 has the greatest contribution to the extracellular secretion of camalexin, our next research focused on whether the kinases OXI1 and AGC2-2 could phosphorylate and regulate the function of PDR6.

Most of the phosphorylation sites of ABC transporters are well conserved among plant or animal orthologs[20]. Given that these sites are mostly contained in the linker region, we performed an *in vitro* kinase assay by using four synthetic peptides contained in intracellular regions (ICRs, **Fig. 6A**) as substrates; these peptides were selected on the basis of *in silico* prediction and phosphor-proteomics[32, 33]. As shown in **Fig. 3A** and **3B**, OXI1 and AGC2-2 reacted with peptides 1, 2, and 4, thus producing a large amount of ADP, whereas neither kinase reacted with peptide 3. Interestingly, the high concentration of peptide 2 inhibited the progress of the reaction possibly due to the inhibition of kinase activity by high substrate concentrations.

**Fig.3.**
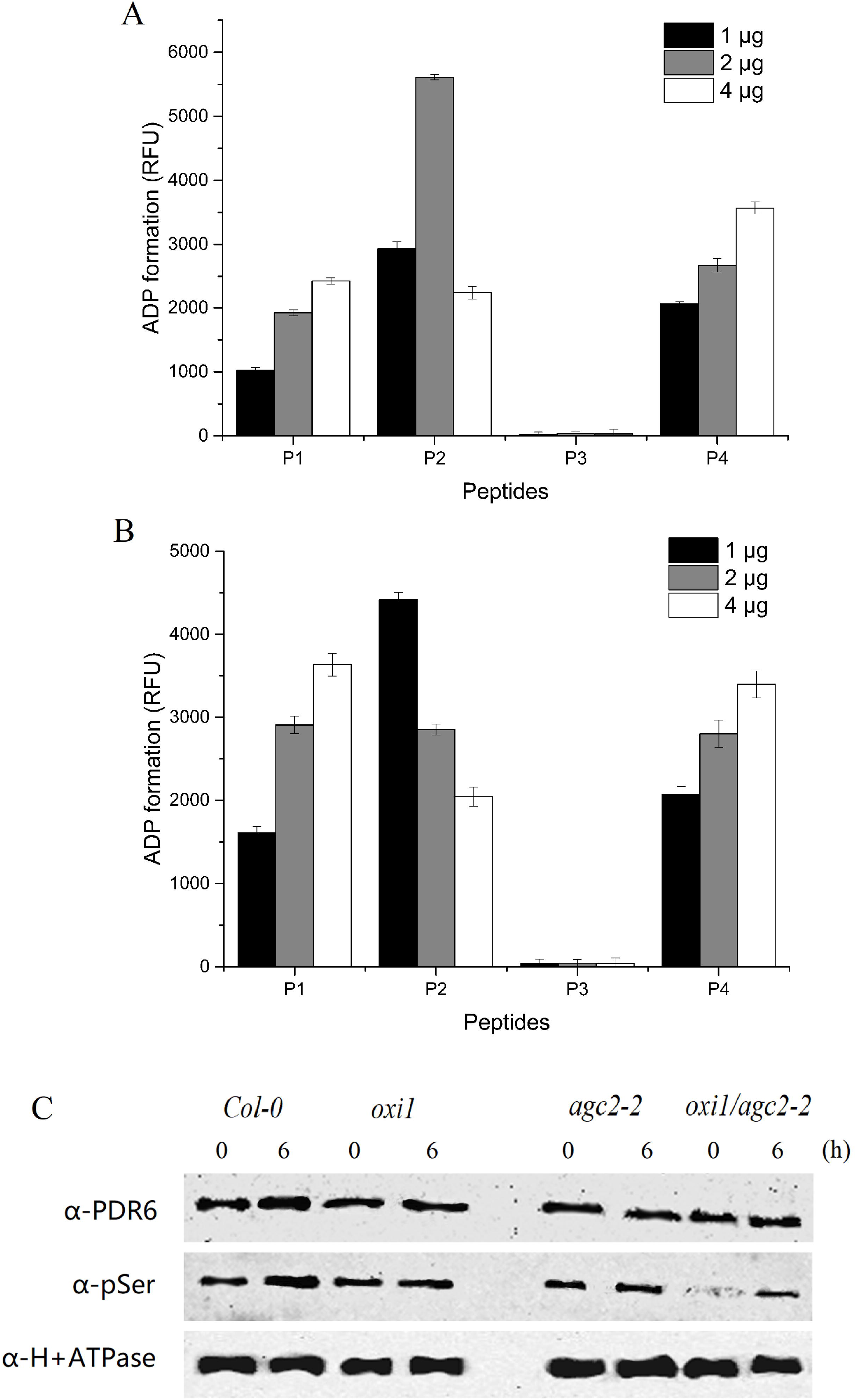
OXI1 and AGC2-2 are related to the phosphorylation of PDR6. Four synthetic peptides of PDR6 located in its ICRs were reacted *in vitro* with *E. coli*-expressed OXI1 (**A**) or AGC2-2 (**B**) by using a universal kinase assay kit (fluorometric, Abcam, USA) in accordance with the manufacturer’s instructions. The fluorescence in wells without the substrate was used as a control and subtracted from the values of the wells with kinase reactions. The *in vivo* phosphorylation levels of PDR6 in Col-0 and the mutants *oxi1, agc2-2*, and *oxi1/agc2-2* were measured through Western blot analysis (**C**). Total plasma membrane proteins were extracted; separated through SDS-PAGE; and immunoanalyzed with anti-PDR6, anti-pSer, and anti-H^+^-ATPase.

We measured the phosphorylation level of PDR6 in wild-type *Arabidopsis* and the two kinase mutants to further explore the effects of the two kinases on the phosphorylation state of PDR6. Chitin induced the expression of PDR6 in Col-0 but showed no significant effects on that in *oxi1, agc2-2*, and *oxi1/agc2-2* (**Fig. 3C**). At 6 h after chitin induction, the phosphorylation level of PDR6 increased significantly in the wild-type and mutant. The lower phosphorylation levels of PDR6 in the *oxi1, agc2-2*, and *oxi1/agc2-2* mutants than those in Col-0 (**Fig. 3C**) illustrated that the two kinases were related to the phosphorylation modification of PDR6. Notably, the phosphorylation of PDR6 in the mutants did not completely disappear. This result indicated that other kinases may be involved.

### OXI1 and AGC2-2 interact with PDR6 *in vitro* and *in vivo*

Protein–protein interactions were examined *in vitro* and *in vivo* to further investigate whether or not OXI1 and AGC2-2 directly phosphorylate PDR6. PDR6 could not be expressed successfully by *Escherichia coli* because it is a protein with multiple transmembrane structure and high molecular weight. Therefore, we expressed and purified only the two ICRs (ICR1 and ICR2) of PDR6 (**Fig. 6A**) by using the classical T7 expression system. The results of the pull-down assay showed that OXI1 and AGC2-2 could interact with the two ICRs (**Fig. 4A**).

**Fig.4.**
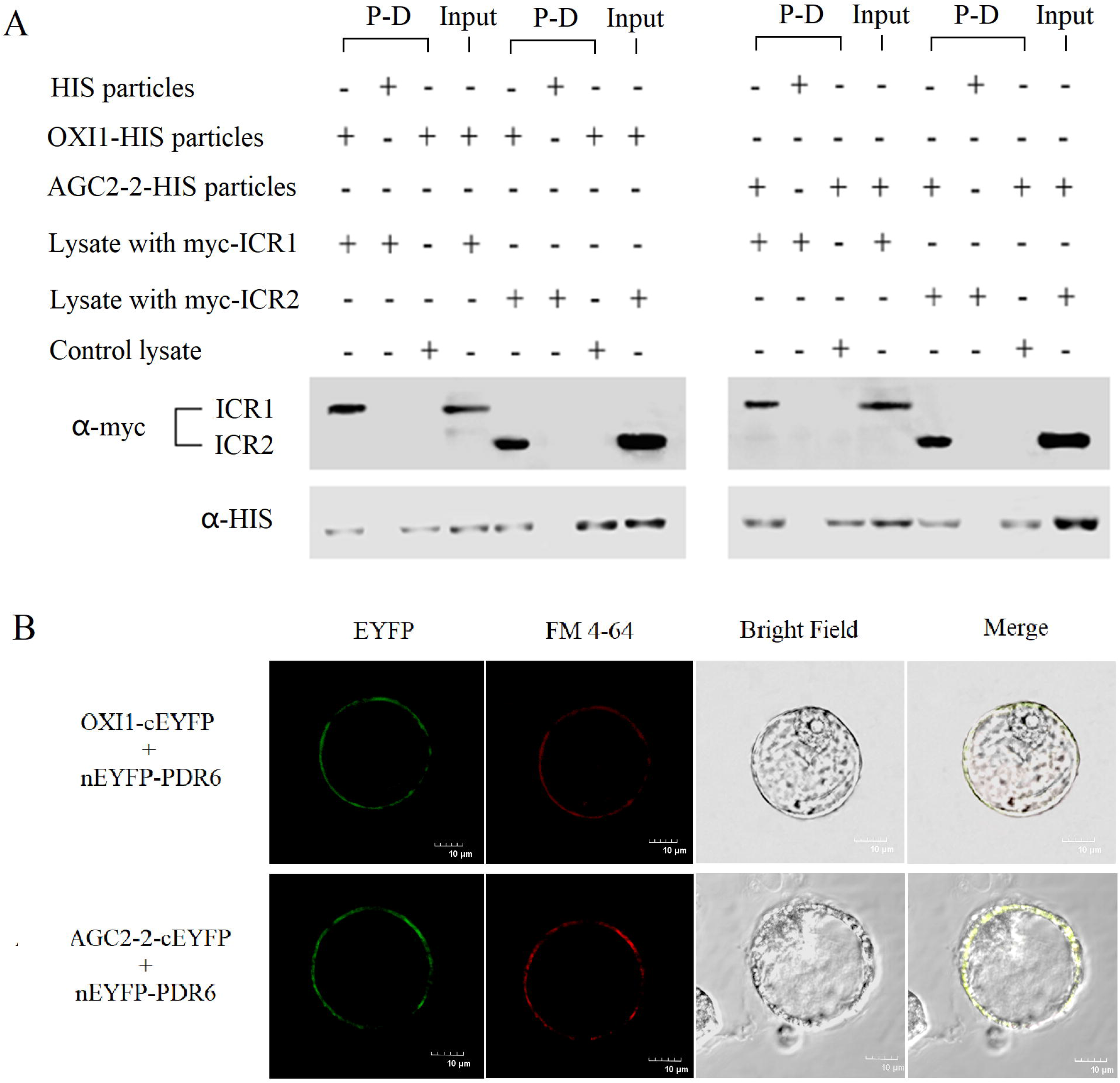
PDR6 directly interacts with the two kinases OXI1 and AGC2-2. For the *in vitro* pull-down assay (**A**), the HIS tag-fused OXI1 or AGC2-2 bound on Ni-particles was incubated with *E. coli* lysate containing either the myc tag-fused ICR1 or ICR2 of PDR6. Empty particles and lysate without IPTG-induced expression (control lysate) were introduced as the controls. For BiFC experiments (**B**), cEYFP-fused OXI1 or AGC2-2 was coexpressed with nEYFP-fused PDR6 in tobacco BY-2 protoplasts *via* electroporation-mediated transformation. FM 4-64 dye was applied to confirm plasma membrane localization. Fluorescence was visualized and captured by using a laser confocal microscope.

We first utilized the bimolecular fluorescence complementation (BiFC) method to verify protein–protein interaction *in vivo*. As shown in **Fig. 4B**, the bright fluorescence on the plasma membrane of tobacco BY-2 protoplasts resulted from the fusion of the C-terminal half of EYFP to OXI1 and that of AGC2-2 complemented with the N-terminal half of EYFP to PDR6. We applied the rapamycin-dependent delocalization method, which is based on the capability of rapamycin to alter the localization of a bait protein and its interactors *via* the heterodimerization of the FKBP and FRB domains[34], to confirm the conclusion inferred from the BiFC results. Given that the FRB domain was anchored at the plasma membrane, after rapamycin infiltration, the FKBP-mCherry-fused ICR1 was delocalized to the plasma membrane from the cytoplasm (**Fig. 5**). When OXI1-EGFP or AGC2-2-EGFP was coexpressed, rapamycin targeted the kinase and the ICR1 of PDR6 to the plasma membrane (**Fig. 5**; ICR2 showed similar results, which are not shown). These results indicated direct contact between the two kinases and ICRs of PDR6 *in vivo*.

**Fig.5.**
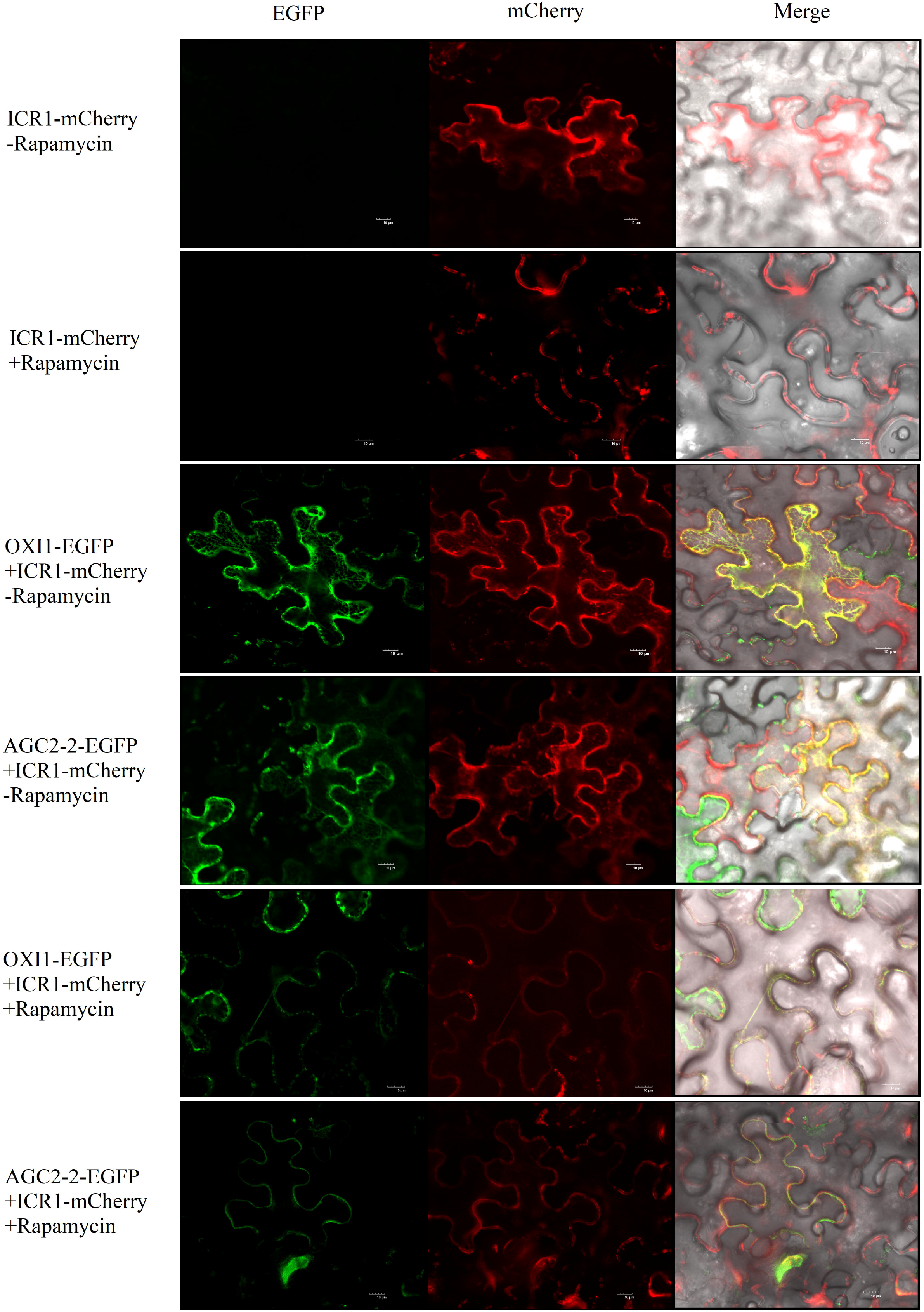
Confirmation of the *in vivo* interactions of the two kinases OXI1 and AGC2-2 with the ICR1 of PDR6 through rapamycin-dependent delocalization. In the presence of rapamycin, ICR1-fused mCherry-FKBP12 was targeted to the plasma membrane of tobacco epidermic cells. The colocalization of mCherry and EGFP, which was fused with OXI1 or AGC2-2, after rapamycin infiltration indicated that an *in vitro* interaction had occurred.

### Identification of the phosphorylation sites of PDR6

The peptide products of the *in vitro* kinase assay were further analyzed through LC–MS/MS to find the target phosphorylation sites of PDR6. Seven phosphorylated serine and threonine sites were identified in three of the four peptides (**Fig. 6A**; MS spectra are shown in **Supplementary File 1**). Among these sites, T11, S31, T42, S827, and T832 were phosphorylated by OXI1; S33, T42, S820, and S827 were phosphorylated by AGC2-2; andT42 and S827 were two common sites shared by both kinases (**Fig. 6B**). Although previously published phosphor-proteomics studies identified many phosphorylation sites in PDR6[35-38], only S33 was identical to our results. Notably, although the phosphorylation sites of the three transporters (PDR6, PDR8, and PDR12) were distributed in conserved linker regions, little conservation was found among the specific sites (**Fig. 6B**).

**Fig.6.**
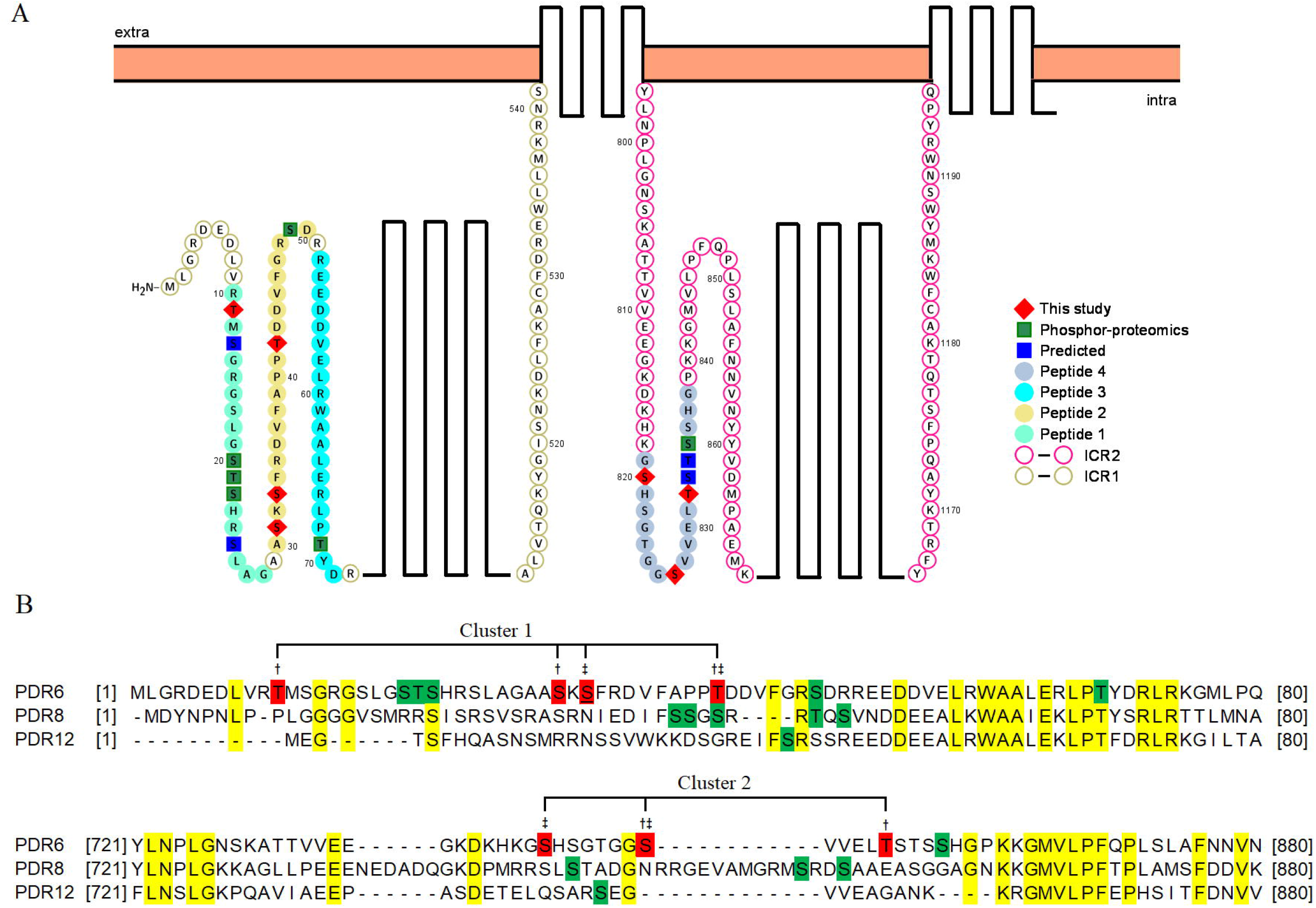
Identification of the phosphorylation sites of PDR6. The locations of ICRs, the four synthetic peptides, and the phosphorylation sites of PDR6 are shown in a schematic of the topological structure of PDR6 drawn by using the Protter web tool[56] (**A**). The phosphorylation sites and conserved amino acids of the three PDR transporters were identified through protein sequence alignment by using MEGA7.0[57] (**B**). † and ‡ indicate the action sites for OXI1 and AGC2-2, respectively. Green highlights: phosphorylation sites identified through phosphor-proteomics; Red highlights: phosphorylation sites identified in this study. The underlined S33 was identified in previous phosphor-proteomics studies and our work.

### Influence of phosphorylation on the function of PDR6

We used site-directed mutagenesis to generate a series of the alanine substitution mutants of the *PDR6*_*pro*_*::PDR6-HIS* construct to mimic the dephosphorylated state and to evaluate the potential roles of the above identified phosphorylation sites in the function of PDR6. Additionally, we created two polymutants containing the phosphorylation sites that were distributed in the two discontinuous linkers (Cluster 1 and Cluster 2, **Fig. 6B**). We transformed each mutant construct into the *pdr6* mutant and assessed the function of each PDR6 variant by evaluating the level of camalexin secretion. The PDR6 variants T11A, T42A, and S820A all were able to fully restore camalexin secretion to levels that were seen in Col-0 and the recomplementary lines that were transformed with *PDR6*_*pro*_*::PDR6-HIS* (**Fig. 7A**). These results showed that phosphorylation at these residues was unrelated to the efflux function of PDR6. The absence of a statistically significant difference in camalexin secretion between the *pdr6* mutant and S31A, S33A, S827A, and T832A (**Fig. 7A**) suggested that these phosphorylation sites are important for the function of the transporter. Intriguingly, the mean camalexin secretion level of the variant S33A was even lower than that of *pdr6*. However, this difference was not statistically significant at the confidence level of 99%. Consistent with the requirement for phosphorylation at serines S31, S33, S827 and T832, the Clusters 1 and 2 polymutant also exhibited impaired efflux function (**Fig. 7A**).

**Fig.7.**
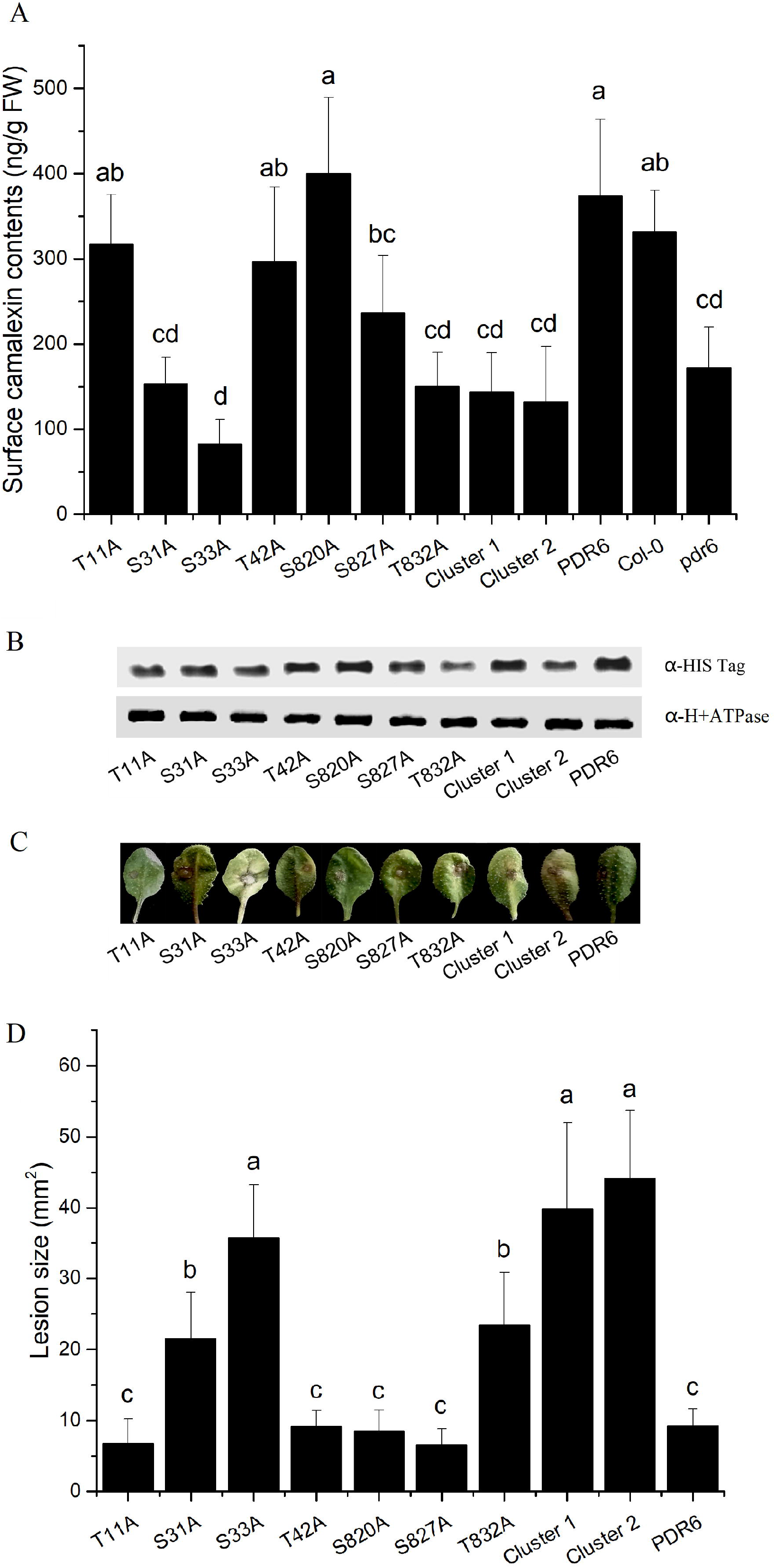
Effects of the dephosphorylation mutation of PDR6 on camalexin secretion, protein contents, and disease resistance to *B. cinerea*. The S-to-A or T-to-A variant expression constructs were expressed in a *pdr6* mutant background driven by a native PDR6 promoter. The secreted camalexin contents were measured after 4 weeks of growth (**A**). The protein contents of the plasma membrane-located PDR6 were examined through Western blot analysis by using anti-HIS antibody, and H^+^-ATPase was set as the internal control (**B**). The resistance to *B. cinerea* was examined, and disease symptoms (**C**) and lesion sizes (**D**) were recorded at 3 days after inoculation. “PDR6” indicates the expression of native PDR6 to recomplement the *pdr6* mutant. Data are presented as mean ± SD, n = 5–6. Letters on scale bars indicate significant differences at p < 0.01 in accordance with Duncan’s multiple range test. OE: overexpression.

Considering that phosphorylation modifications may change the stability or subcellular localization of the transporter, we next examined the abundance of the PDR6 protein in the plasma membrane of the transgenic variants. Our results showed that the variant S832A and the polymutant Cluster 2 had reduced PDR6 protein levels in the plasma membrane (**Fig. 7B**). This result demonstrated that the S832 amino acid might be related to the stability of PDR6. The consistency of the disease resistance of PDR6 variants to *B. cinerea* with the extracellular contents of camalexin (**Fig. 7C–D**) further confirmed the function of phosphorylation sites in resistance responses.

## Discussion

Transporter phosphorylation modification may affect transporter activity, stability, and/or subcellular localization[14]. In plant breeding research, the functional manipulation of secondary metabolite transporter(s) can confer crops with strengthened disease and insect resistance[39]. Therefore, the study of the phosphorylation modification of secondary metabolite transporters has important research value. Although previous studies have reported that the transporters PDR6, PDR8, and PDR12 are related to the extracellular secretion of camalexin in *Arabidopsis*, their conclusions have been inconsistent[12, 13]. This study verified the activity of the three transporters and found that PDR6 plays a major role in transporting camalexin. The results of this study support the findings of Khare et al. The lack of consensus among studies may be caused by the different methods used for measuring camalexin contents; for example, He et al. applied the water immersion method, whereas we and Khare et al. utilized the *n*-hexane extraction method. The significantly lower total content of camalexin detected in this study than that found in the above two studies may be due to different culture conditions considering that some cases with moderate or low levels of camalexin have also been reported[40, 41].

By detecting the contents of camalexin on the leaf surfaces of mutants and overexpression lines, we confirmed that the two kinases OXI1 and AGC2-2 are related to the extracellular secretion of camalexin. The overexpression of the two kinases increased the camalexin content on the leaf surface and decreased that inside the leaf, thereby enhancing the resistance to *P. syringae* and *B. cinerea* inoculated through a noninjection method. However, Petersen et al.[24] found that the overexpression of *OXI1* reduced the resistance of *Arabidopsis* to *P. syrengae* and that the expression of *OXI1* was not correlated with resistance to *B. cinerea*. Given that camalexin-resistant and -sensitive isolates can be found within one pathogen species as it was shown for *B. cinerea*[42], the inconsistency of the results may be attributed to differences in pathogenic capacity between different strains. We speculated that the results for *P. syringae* differed due to the inconsistency of the inoculation methods used as discussed in the Results section. Petersen et al. also tested the pathogenicity of an oomycete pathogen *H. parasitica* through leaf surface inoculation and found that *OXI1* overexpression was also not conducive to the resistance to this pathogen. Although we cannot provide a reasonable explanation for this phenomenon, we can speculate that OXI1 weakens the resistance to the pathogen because it also affects other defense responses while regulating the distribution of camalexin. In rice, the overexpression of *OsOxi1*, an AGC kinase homologous to OXI1, enhances basal resistance to the blast fungus *Magnaporthe oryzae*, indicating that OsOxi1 positively regulates disease resistance[43]. This result is consistent with our findings.

In addition to being associated with pathogen resistance, OXI1 is involved in interactions among beneficial microbes, pests, and their host plants. Camehl et al.[26] showed that the normal OXI1 gene is necessary for the endophytic fungus *Piriformospora indica* to promote the growth of *Arabidopsis*. The plant growth-promoting effects of multiple bacterial strains need root-specific camalexin biosynthesis[44]. Therefore, OXI1 may facilitate the interaction between beneficial microorganisms and host plants by increasing camalexin secretion. However, an experiment with pests showed that the presence of OXI1 is detrimental to the resistance of *Arabidopsis* to *Myzus persicae*[45]. The OXI1-mediated reduction of intracellular camalexin content may be one of the reasons for the improved reproduction of piercing-sucking aphids because camalexin synthesis is also required for controlling the green peach aphid and cabbage aphid infestation of *Arabidopsis*[46].

Notably, OXI1 and AGC2-2 are likely to be involved in the extracellular secretion of other compounds in addition to the distribution of camalexin. For example, when using HPLC to analyze the metabolite content on the leaf surface, a substance with a retention time of 11.75 min was found to have a significantly higher content in Col-0 than in the two mutants (**Fig. 1B**). However, the identification of this compound still awaits further research.

Considering that the overexpression of *OXI1* and *AGC2-2* did not lead to changes in the transcriptional levels of the three transporters, we speculated that the two kinases mediated the changes in camalexin distribution in *Arabidopsis* due to the direct or indirect phosphorylation modification of the transport proteins. This modification, in turn affected, their activity. Although subsequent studies revolved around PDR6 and confirmed that OXI1 and AGC2-2 can directly phosphorylate PDR6, additional *in vitro* kinase assays indicated that PDR8 and PDR12 are also likely to act as the substrates of kinases (**Fig. S3**).

In *Arabidopsis* and rice, chitin is recognized by CHITIN ELICITOR RECEPTOR KINASE 1, which then induces an elevation in ROS levels[47]. OXI1 is required for the full activation of MAPK3 and MAPK6 under H_2_O_2_ stress[23]. In addition to H_2_O_2_, 3-phosphinositide-dependent protein kinase 1 downstream of the phosphatidic acid signal mediates the phosphorylation of OXI1[48]. Following studies have shown that PTI family members are phosphorylated by OXI1 and mediate the regulation of MAPKs by OXI1[25, 49]. In this study, chitin was shown to induce PDR6 phosphorylation, which was mainly dependent on OXI1 and AGC2-2 (**Fig. 3C**). Therefore, we speculated that the recognition of chitin and the generation of ROS increase the phosphorylation levels of OXI1, which in turn phosphorylates PDR6 and enhances its function. OXI1 may also phosphorylate other PDR transporters at the same time, thereby regulating the distribution of other metabolites in *Arabidopsis*.

We first identified seven potential phosphorylation sites through LC–MS/MS and then examined their function *via* dephosphorylation mutations to verify whether OXI1 can enhance the function of PDR6 through phosphorylation modification. The experimental results confirmed that the action site S31 of OXI1 and the action sites S33 and S827 of AGC2-2 had positive effects on the efflux activity of PDR6. In addition, the action sites of OXI1, T832, may contribute to the plasma membrane localization of PDR6 or enhance the stability of PDR6. Although the phosphorylation sites identified through *in vitro* kinase reactions are not completely identical to those previously identified through phosphoproteomics, some of these phosphoproteomic results are based on various treatments, such as auxin irritation[36] and nitrogen deprivation[37]. Under different environments and stresses, the phosphorylation modification of a specific ABC transporter may occur at different positions, thereby showing different effects. For example, in contrast to the multiple sites identified through proteome analysis, only one phosphorylation site that was related to the upstream kinase PID was identified in ABCB1[20].

In conclusion, this study discovered for the first time the upstream kinases acting on plant PDR transporters that efflux secondary metabolites. The direct phosphorylation of PDR6 by OXI1 and AGC2-2 regulated the efflux function of PDR6 and the disease resistance of *Arabidopsis*. However, whether OXI1 and AGC2-2 can directly regulate the function of PDR8 and PDR12 and whether they affect the distribution of other secondary metabolites remain to be further investigated. OXI1 and AGC2-2 may not be the only kinases targeting PDR6. Given that PDR transporters are often shown to be able to transport several structurally unrelated compounds and participate in a very diverse range of physiological processes[8], studying the interrelationships between different kinases targeting one PDR transporter and their phosphorylation sites will be interesting.

## Materials and Methods

### Plant materials and growth

The mutants *pdr6* (SAIL_5_G10)[12], *pdr8* (SALK_110926; Stein et al., 2006), *pdr12* (SALK_005635; Campbell et al., 2003), *oxi1* (CS2103498; Bolle, Huep et al. 2013), *agc2-2* (CS2103499; Bolle, Huep et al. 2013), and *oxi1/agc2-2* (CS2103497;[30]) were all obtained from the Arabidopsis Biological Resource Center.

The seeds of all the plant materials were germinated in 0.5× Murashige Skoog medium[50], 1% sucrose, and 0.7% agar. The seeds were stratified at 4 °C for 72 h, germinated at 22 °C under continuous day conditions, and then transferred into small pots for growth for 4 weeks in controlled environment chambers under a 14:10 h light/dark cycle at 22 °C and 55% relative humidity.

### Generation of transgenic plants

Each CDS without the stop codon was PCR-amplified from a cDNA library of *Arabidopsis* Col-0 and ligated into the *Hind*III and *EcoR*I restriction sites of the modified plant binary expression vector pYBA1132-35SN (**Fig. S6A**) to construct 35S_pro_::OXI1 and 35S_pro_::AGC2-2 expression plasmids. For the construction of the PDR6_pro_::PDR6 plasmid, 2166 nucleotides upstream of the start codon (NC_003071.7: 15 255 418 to 15 257 583) were artificially synthesized (GeneralBiol, China) and ligated into the *Age*I and *BamH*I sites of pYBA1132-35SN. The CDS of PDR6 without the stop codon was amplified and ligated into the pUC57 cloning plasmid and subcloned into the *Hind*III and *EcoR*I sites via recombination. The phosphorylation sites were mutated into Ala through site-directed mutagenesis.

The vectors were individually transformed into *Agrobacterium tumefaciens* (strain GV3101), and flowering Col-0 or *pdr6* plants were dipped into *A. tumefaciens*-containing infiltration medium as previously described[51]. T1 seeds were surface-disinfected and plated on Murashige and Skoog agar plates containing 50 μg kanamycin/mL. After the confirmation of positive lines through quantitative RT-PCR or Western blot analysis with the anti-HIS antibody, 3 to 5 lines were selected for further experiments.

### Plant treatment and inoculation

Four-week-old plants were sprayed with chito-octaose (Toronto Research Chemicals, Canada) at the concentration of 2 μM or with distilled water. After the indicated treatment time, whole rosettes were dissected and treated for analysis immediately.

For the inoculation of the virulent strain *P. syringae* pv. *tomato* DC 3000, the entire rosette was dipped into the bacterial suspension (1 × 10^8^ CFU/mL in water supplemented with 0.02% Silwet L-77) for 5 s. After 3 days of treatment, the colony counting method was used to analyze the bacterial number on leaf disks[52]. For the inoculation of *B. cinerea*, 5 μL of the spore suspension (5 × 10^5^ spores/mL in 1% Sabouraud maltose broth buffer) was dropped onto detached leaves. Inoculated leaves were kept under high humidity for 3 days before lesion size measurement.

### Camalexin content determination

The rosettes of three plants were combined and immersed into 30 mL of *n*-hexane to measure the amounts of camalexin secreted on the leaf surface. After 30 s of extraction with gentle shaking, the *n*-hexane extracts were subjected to rotary evaporation under vacuum and redissolved in 200 μL of methanol before HPLC–UV analysis. After *n*-hexane extraction, the rosettes were further used for the determination of intracellular camalexin contents, and the samples were prepared in accordance with the method of Beets and Dubery[53]. The HPLC–UV method was performed in reference to the protocol of Beets and Dubery[53] except for the use of a Waters 1525 binary pump with Waters 2998 photodiode array detector and a Waters Sunfire C18 (150 × 4.6 mm, 5 μm) column.

### *In vitro* kinase assay

The CDS of *OXI1* and *AGC2-2* was cloned into pET-28a(+) to obtain the purified kinases. His-tagged OXI1 and AGC2-2 fusion proteins were expressed in *E. coli* BL21 (DE3) through IPTG induction. Proteins were purified by using MagneHis™ Protein Purification System (Promega, USA). The peptides of the transporters PDR6, PDR8, and PDR12 were synthesized as substrates by GenScript Biotech (China). For the *in vitro* kinase assay, a universal kinase assay kit (fluorometric) (Abcam, USA) was applied in accordance with the manufacturer’s instructions. Briefly, 20 µL of the ADP assay buffer; 20 µL of the ADP sensor buffer; 10 µL of the ADP sensor; 1 µM ATP; 0.5 µg of purified OXI1 or AGC2-2; and 1, 2, or 4 μg of synthetic peptide as the substrate were added into each tube to a total ADP assay volume of 50 µL/sample. The reaction mixture was incubated at room temperature for 30 min. Fluorescence intensity was measured at the excitation/emission of 540/590 nm in black plates with a SpectraMax M2e microplate reader (Molecular Devices). The fluorescence in wells without the addition of the substrate was used as the control and subtracted from the values for those wells with the kinase reactions.

### *In vitro* pull-down assay

The DNA sequences of ICR1 and ICR2 were fused with the myc tag at their 5D termini through PCR and cloned into pET-11b for expression in BL21(DE3). One milliliter of recombinant *E. coli* containing myc-ICR1 or myc-ICR2 was lysed in buffer (20 mM Tris-HCl, pH 7.5, 100 mM NaCl, and protease inhibitor cocktail). The supernatant was recovered after centrifugation at 14 000 × *g* for 15 min at 4 °C. Then, this lysate was incubated with MagZ™ particle-bound OXI1-HIS or AGC2-2-HIS for 1 h at room temperature. Lysate obtained under non-induced condition was used as the control lysate. Following incubation, the particles were collected in a strong magnetic field and washed with buffer (20 mM 1,3-diazole, 20 mM Tris-HCl, pH 7.5) three times. The bound proteins were eluted from the particles with an elution buffer (300 mM NaCl, 500 mM 1,3-diazole, 20 mM Tris-Cl, pH 7.5) and further analyzed through Western blot analysis with anti-myc (Abcam, USA) and anti-HIS antibodies (Santa Cruz Biotechnology, USA).

### *In vivo* protein interaction

For the BiFC experiment, the CDS regions of OXI1, AGC2-2, and PDR6 were cloned into the expression plasmids pSAT4A-cEYFP-N1 and pSAT6-nEYFP-C1[54], respectively. Tobacco BY-2 protoplast preparation and transient expression *via* electroporation were conducted in accordance with the method of Miao and Jiang[55]. Fluorescence was visualized and captured by using a laser confocal microscope.

For rapamycin-dependent delocalization, a DNA fragment containing two expression cassettes was synthesized artificially (GenScript, China): one was for the plasma-membrane-targeted mTagBFP2-FRB and the other was for the ICR1- or ICR2-fused mCherry-FKBP12 (**Fig. S6B**). The CDS of OXI1 or AGC2-2 was introduced into the pYBA1132 plasmid through BamHI and XhoI restriction cutting and ligation to obtain the EGFP-fused recombinant binary expression plasmid.

These plasmids were transformed into the leaves of *Nicotiana benthamiana* through *A. tumefaciens*-mediated transformation. After 2 days of transformation, the fluorescence in leaves was observed under a laser confocal microscope before and after 2 h of infiltration with 1 μM rapamycin.

### Identification of phosphorylation sites

A total of 90 μL of 50 mM NH_4_HCO_3_ solution was added to 10 μL of the products of the *in vitro* kinase assay. After reduction with 10 mM DTT at 56 °C for 1 h and alkylation with 20 mM IAA at room temperature in the dark for 1 h, the extracted peptides were lyophilized to near dryness and resuspended in 10 μL of 0.1% formic acid before LC–MS/MS analysis.

An Ultimate 3000 system (ThermoFisher Scientific, USA) equipped with 100 μm × 10 cm in-house-made column packed with a reverse-phase ReproSil-Pur C18-AQ resin (3 μm, 120 Å, Dr. Maisch GmbH, Germany) was used for separation. Mobile phase A was 0.1% formic acid in water, and mobile phase B was 0.1% formic acid in acetonitrile. The LC linear gradient was set as follows: from 6% to 9% B for 5 min, from 9% to 50% B for 50 min, from 50% to 95% B for 3 min, and from 95% to 95% B for 2 min. A total of 5 µL of the sample was loaded with the flow rate of 300 nL/min.

Q Exactive™ Hybrid Quadrupole-Orbitrap™ Mass Spectrometer (Thermo Fisher Scientific, USA) was used for fragmentation (spray voltage: 2.2 kV; capillary temperature: 270 °C) and mass analysis (activation type: CID; min. signal required: 1500.0; isolation width: 3.00; normalized coll. energy: 40.0; default charge state: 6; activation Q: 0.250; activation time: 30.000; MS precursor m/z range: 50.0–1500.0; data-dependent MS/MS: up to the top 15 most intense peptide ions from the preview scan in Orbitrap). The obtained source spectrum was input into Maxquant (1.6.2.10) for the identification of peptide and phosphorylation sites.

### Western blot analysis

For the isolation of plasma membrane proteins, Minute™ Plant Plasma Membrane Protein Isolation Kit (Invent Biotechnologies, USA) was used in accordance with the manufacturer’s instructions. Proteins were separated on 5%–12% SDS-PAGE gels. For the detection of PDR6 and its phosphorylation level, the gels were firstly analyzed through immunoblotting with anti-PDR6 antibody (prepared with peptide CTVVEEGKDKHKGSH as antigen by GenScript, China) and a horseradish peroxidase-conjugated antirabbit IgG secondary antibody (Beyotime Biotechnology, China). The membrane was stripped in buffer (15 g/L glycine, 1 g/L SDS, 10% Tween-20, pH 2.2) for two times and applied for the immunodetection of phosphorylation levels by using an anti-pSer primary antibody (Santa Cruz, USA) and antimouse secondary antibody (Santa Cruz, USA). For each sample, the amount of plasma membrane H^+^-ATPase was analyzed in parallel to eliminate the error caused by different sample loadings by using an anti-H^+^-ATPase antibody (Agrisera, Swedish) and a horseradish peroxidase-conjugated antirabbit IgG secondary antibody (Beyotime Biotechnology, China). The horseradish peroxidase reaction was developed with an enhanced chemiluminescence system (Beyotime Biotechnology, China).

## Accession Numbers

PDR6/ABCG34: AT2G36380; PDR8/PEN3/ABCG36: AT1G59870; PDR12/ABCG40: AT1G15520; OXI1/AGC2-1: AT3G25250; AGC2-2: AT4G13000.

## Figure captions

**Supplementary File 1** MS spectra of the phosphorylation sites identified through LC–MS/MS.

**Fig. S1** Morphology of the gene-overexpressing lines of OXI1 and AGC2-2 was similar to that of Col-0.

**Fig. S2** Contents of camalexin on the leaf surfaces of the mutants *pdr6, pdr8*, and *pdr12*. The sign “asterisk” indicates significant differences between groups at p < 0.05 (one asterisk) or p < 0.01 (two asterisks) in accordance with one-way ANOVA.

**Fig. S3** Peptides from the ICR of PDR8 and PDR12 (**A**) reacted with purified OXI1 or AGC2-2 *in vitro* (**B**).

**Fig. S4** Expression structures of pYBA1132-35SN constructed in our lab (**A**) and pYBA1132-PM-FRB-FKBP12-ICR used for rapamycin-dependent delocalization experiments (**B**).

