## Supplementary Data for "PDR6-mediated camalexin efflux and disease resistance are regulated through direct phosphorylation by the kinases OXI1 and AGC2-2"

**Fig. S1** Morphology of the gene-overexpressing lines of *OXI1* and *AGC2-2* was similar to that of Col-0.

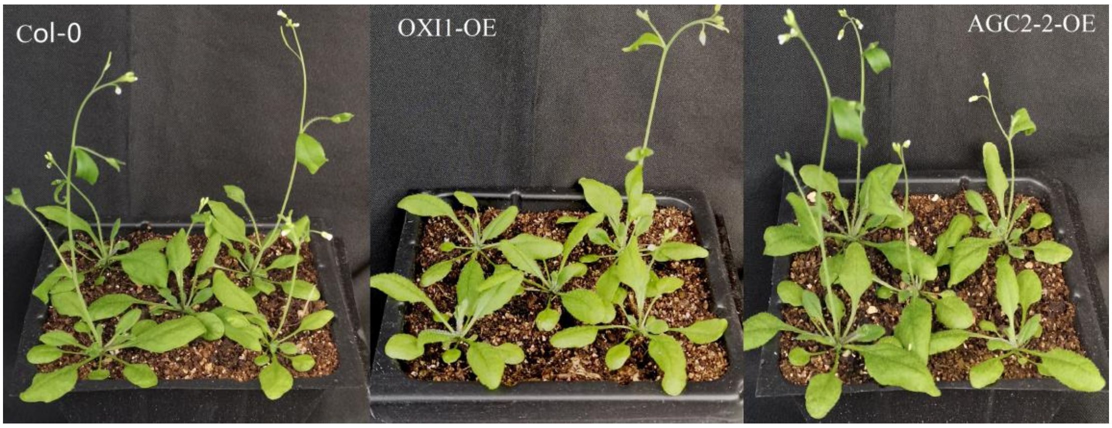

**Fig. S2** Contents of camalexin on the leaf surfaces of the mutants *pdr6*, *pdr8*, and *pdr12*. The sign "asterisk" indicates significant differences between groups at $p < 0.05$ (one asterisk) or $p < 0.01$ (two asterisks) in accordance with one-way ANOVA.

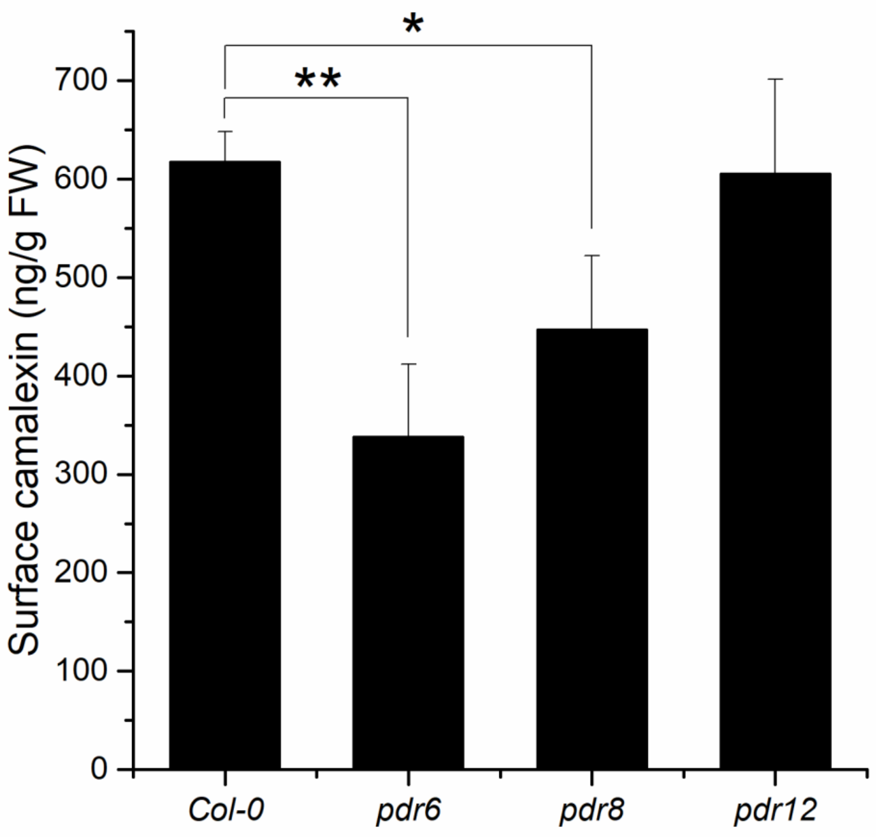

**Fig. S3** Peptides from the ICR of PDR8 and PDR12 (**A**) reacted with purified OXI1 or AGC2-2 *in vitro* (**B**).

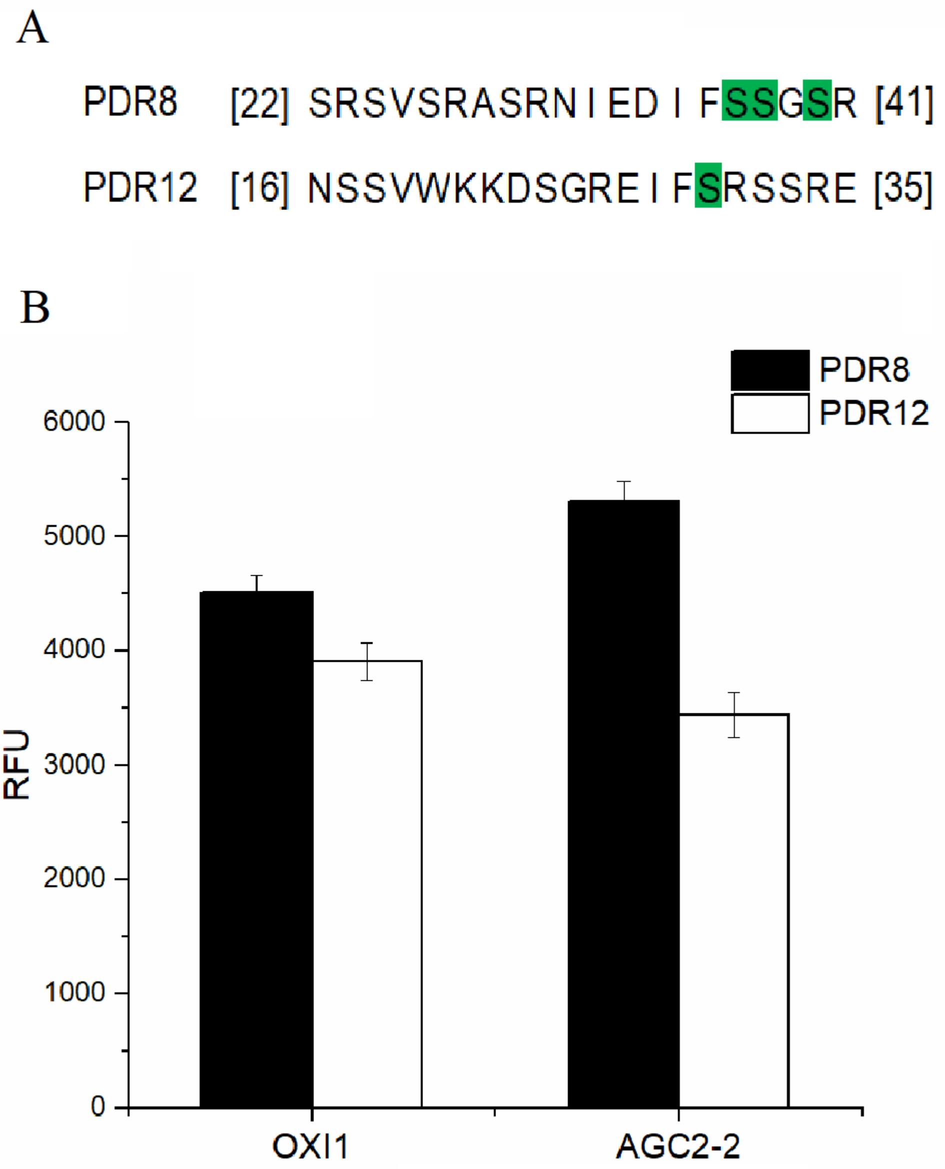

**Fig. S4** Expression structures of pYBA1132-35SN constructed in our lab (**A**) and pYBA1132-PM-FRB-FKBP12-ICR used for rapamycin-dependent delocalization experiments (**B**).

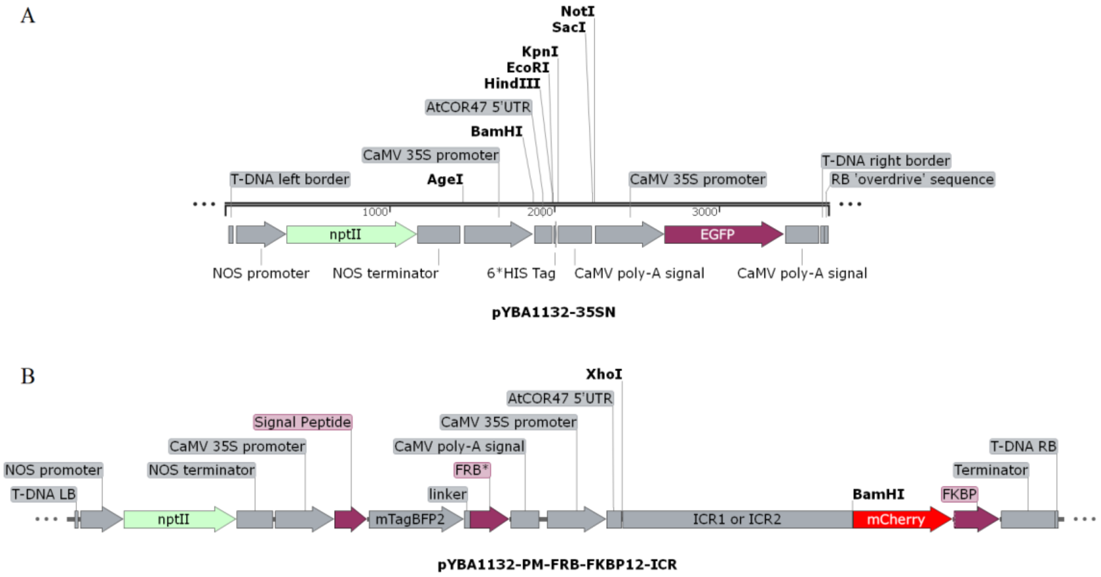

**Supplementary File 1** MS spectra of the phosphorylation sites identified through LC–MS/MS.

pT11 by OXI1

| Raw File | Scan | Method | Score | m/z |
| --- | --- | --- | --- | --- |
| P1_AGC2_1 | 9969 | FTMS; HCD | 49.99 | 500.49 |

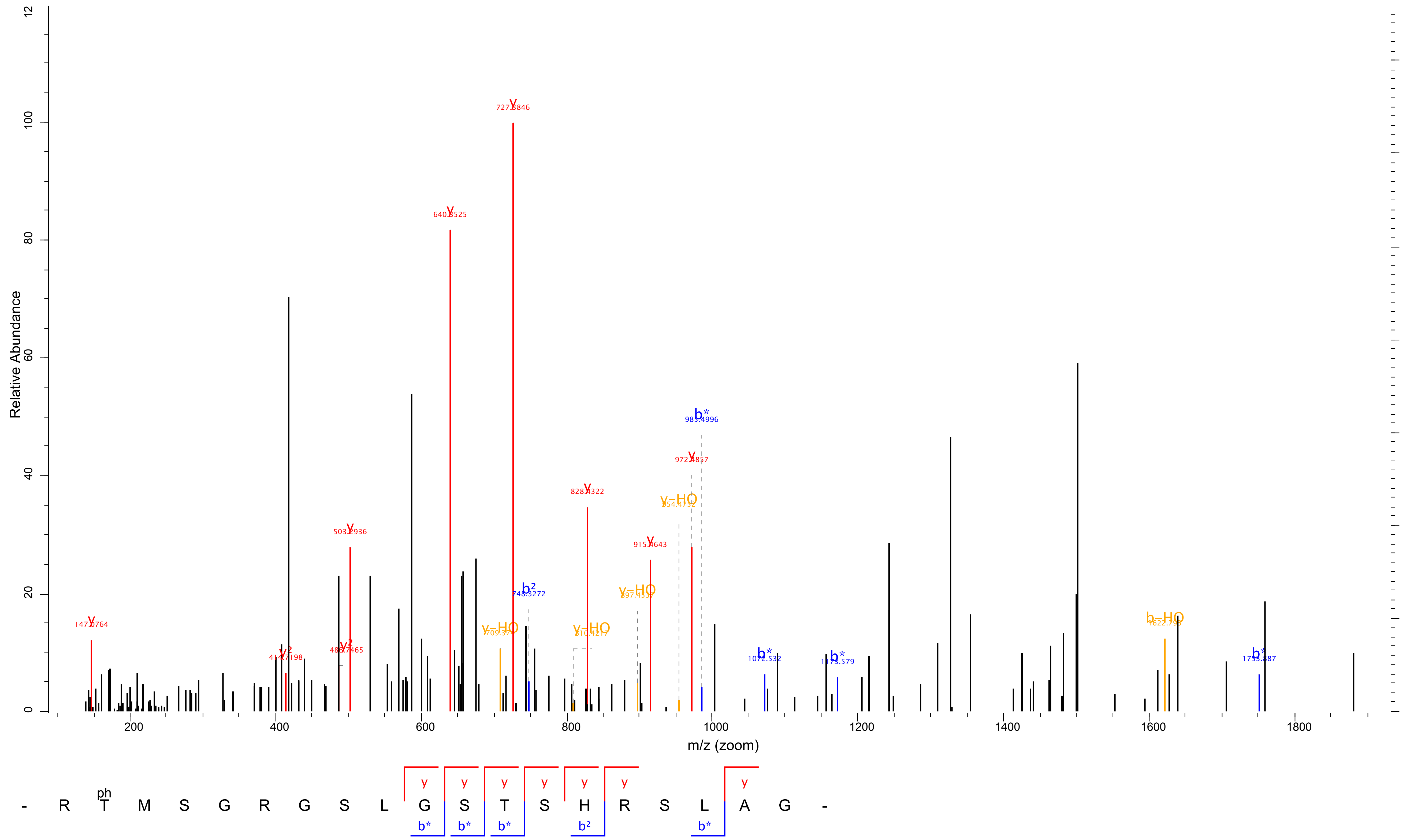

pS31 by OXI1

| Raw File | Scan | Method | Score | m/z |
|---|---|---|---|---|
| P2_2_1 | 21724 | FTMS; HCD | 95.5 | 1197.54 |

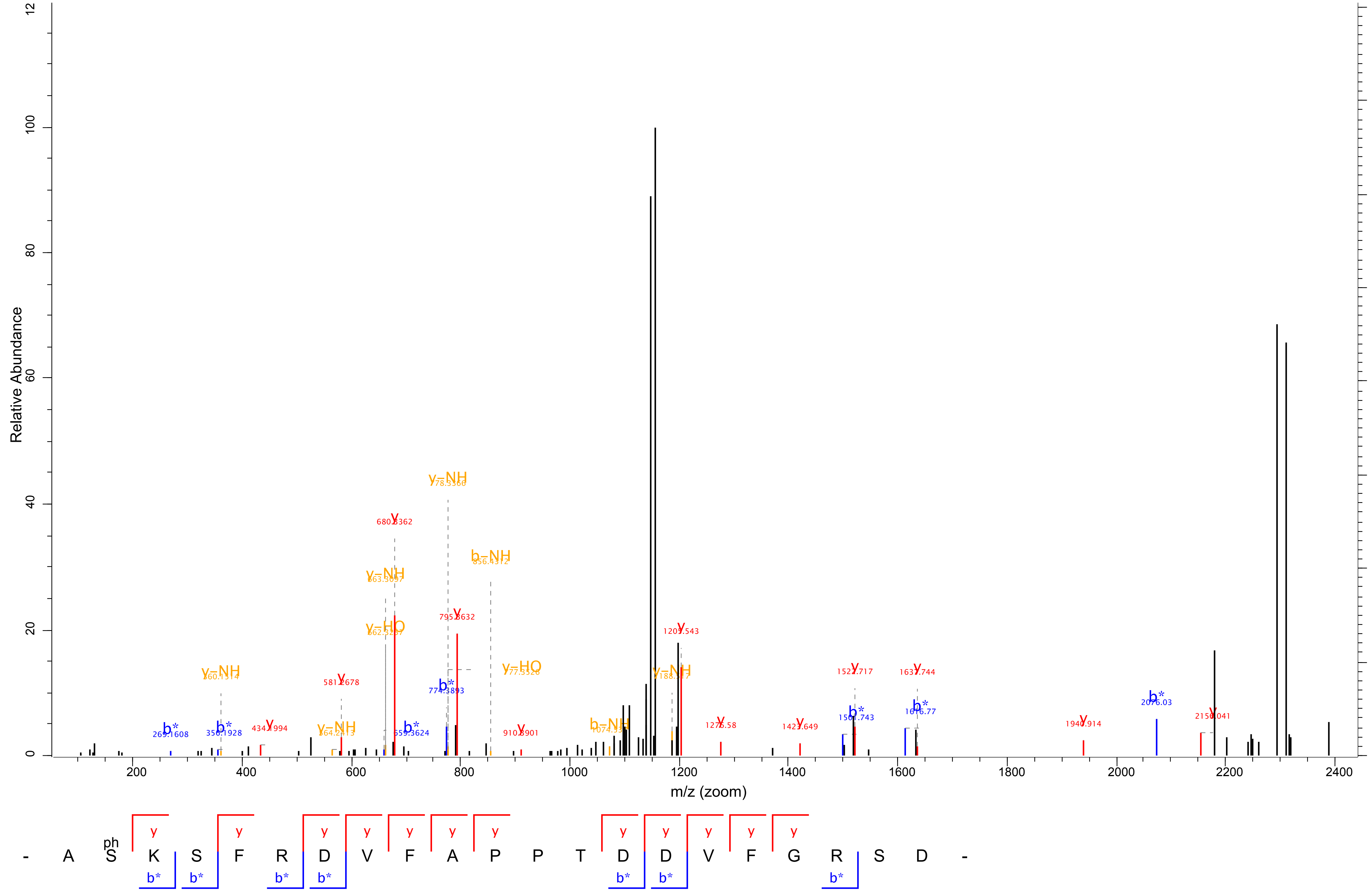

| Raw File | Scan | Method | Score | m/z |
|---|---|---|---|---|
| P2_2_1 | 21801 | FTMS; HCD | 97.63 | 1198.04 |

### pT42 by OXI1

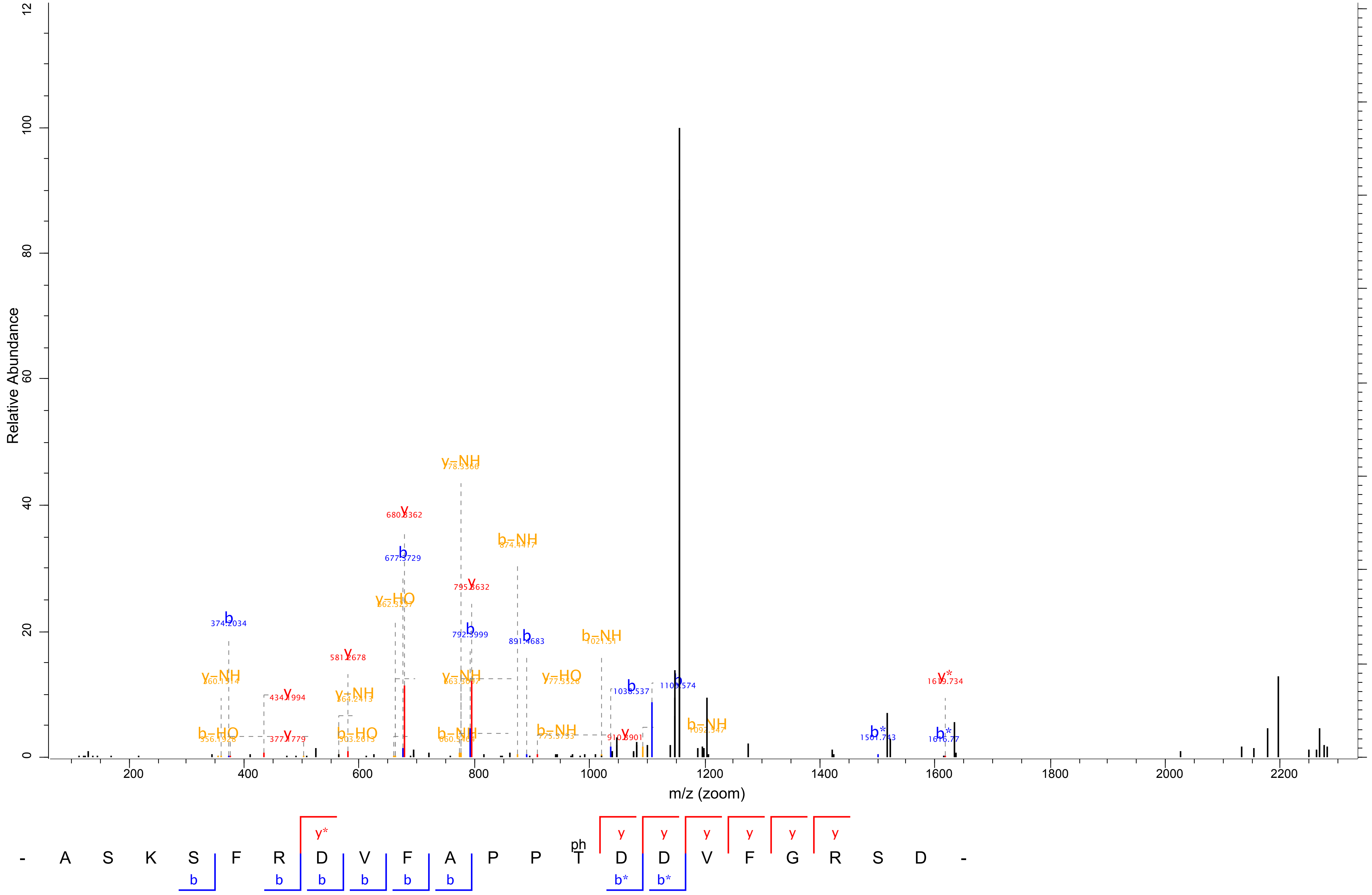

pS827 by OXI1

| Raw File | Scan | Method | Score | m/z |
|---|---|---|---|---|
| P3_2_1 | 4236 | FTMS; HCD | 96.6 | 962.41 |

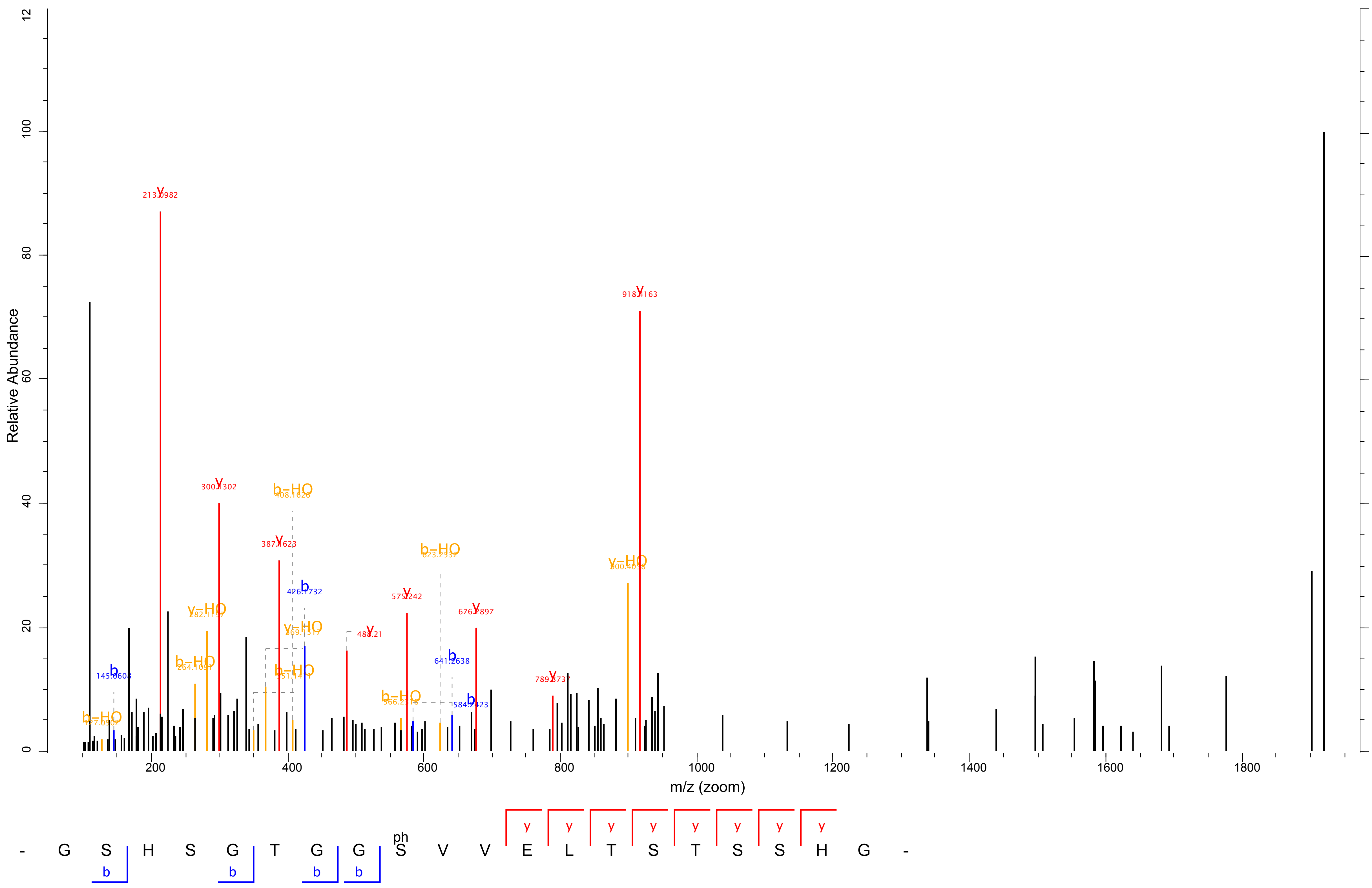

| Raw File | Scan | Method | Score | m/z |
|---|---|---|---|---|
| P3_2_1 | 6757 | FTMS; HCD | 119.76 | 962.92 |

pT832 by OXI1

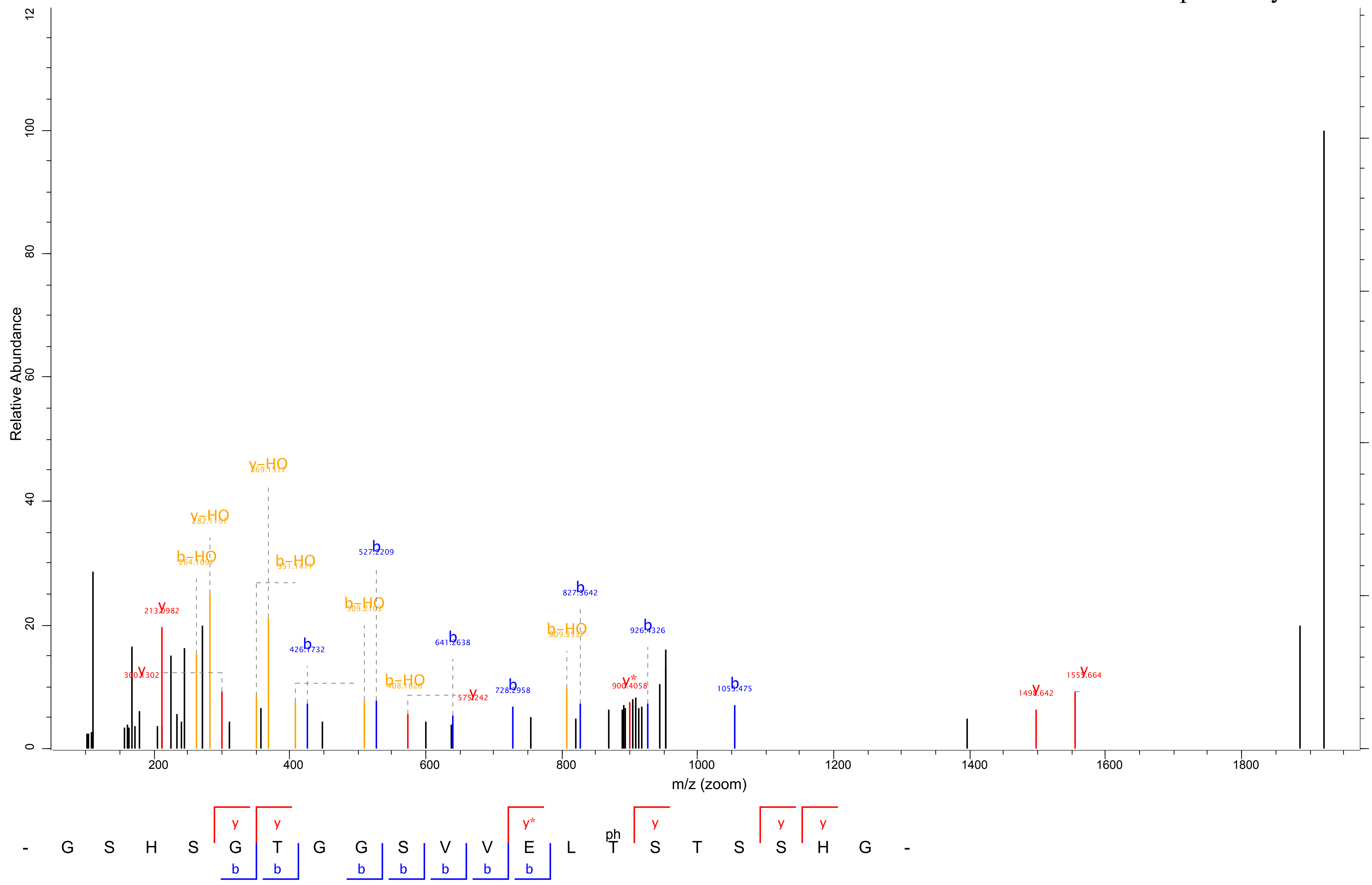

| Raw File | Scan | Method | Score | m/z |
|---|---|---|---|---|
| P2_2_2 | 15038 | FTMS; HCD | 91.62 | 799.04 |

### pS33 by AGC2-2

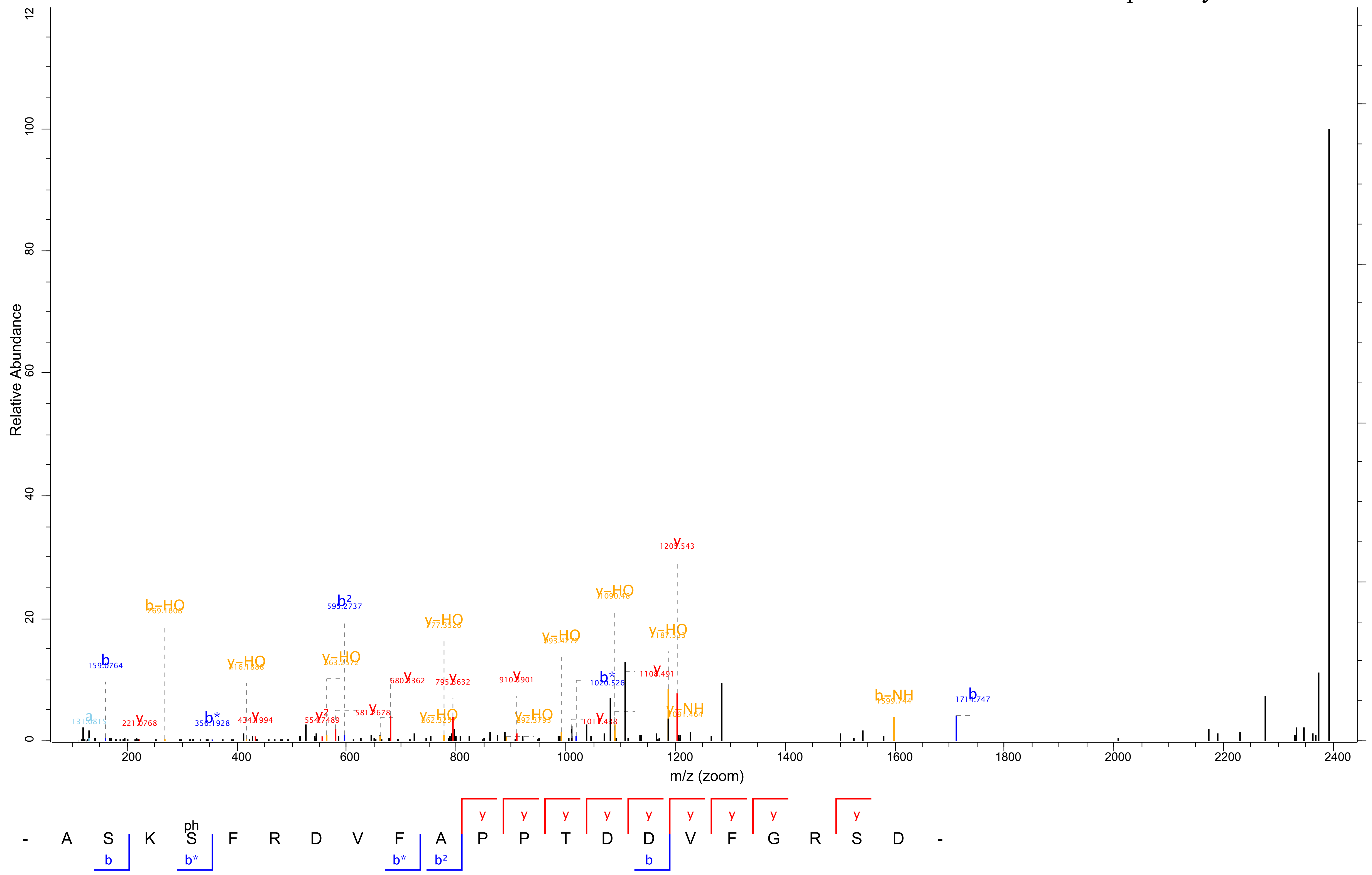

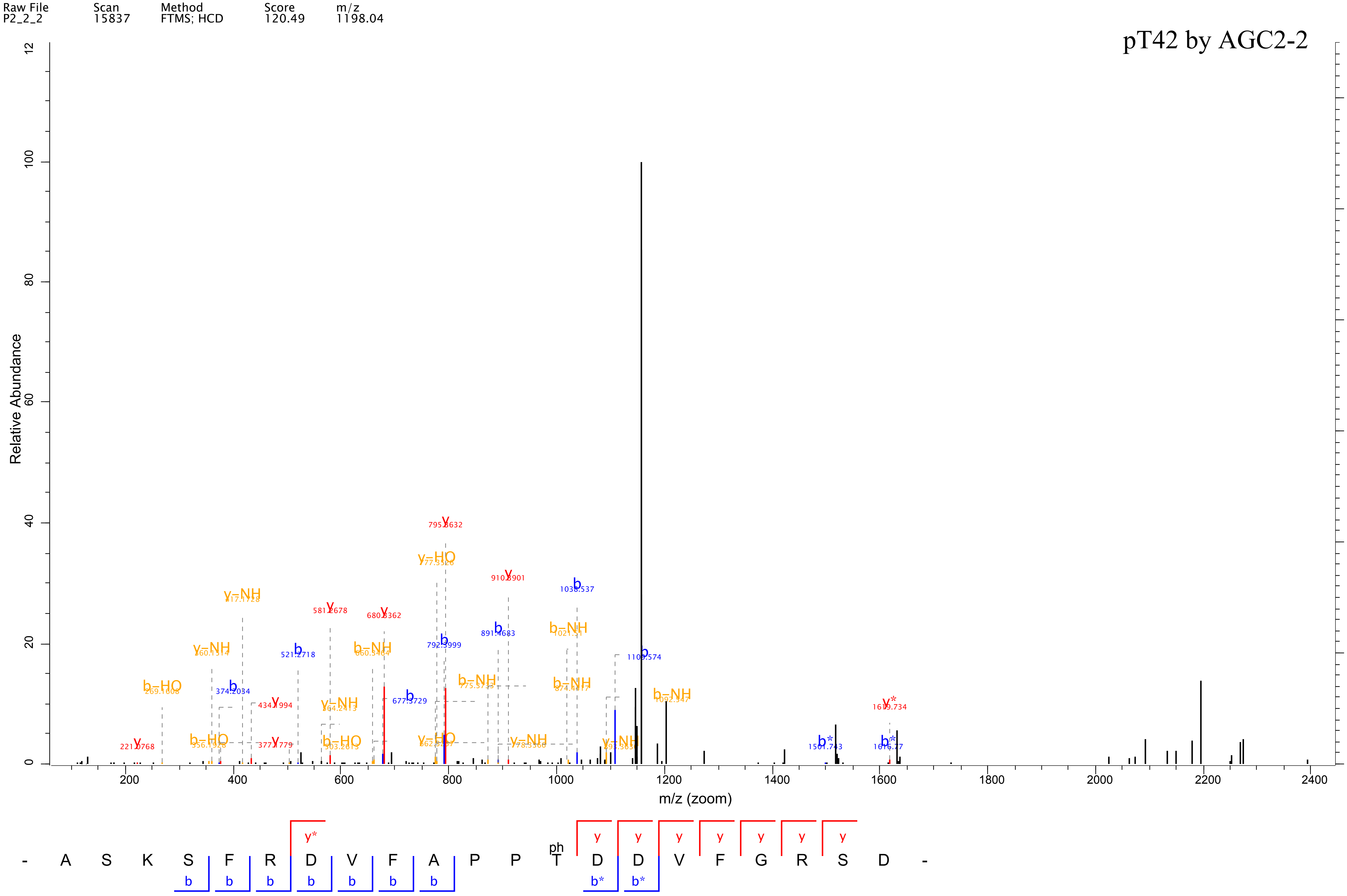

Raw File P2_2_2
Scan 15837
Method FTMS; HCD
Score 120.49
m/z 1198.04
pT42 by AGC2-2
Relative Abundance
m/z (zoom)
- A S K S F R D V F A P P T D D V F G R S D -
ph

| Raw File | Scan | Method | Score | m/z |
| --- | --- | --- | --- | --- |
| P3_2_2 | 5429 | FTMS; HCD | 332.02 | 962.91 |

### pS820 by AGC2-2

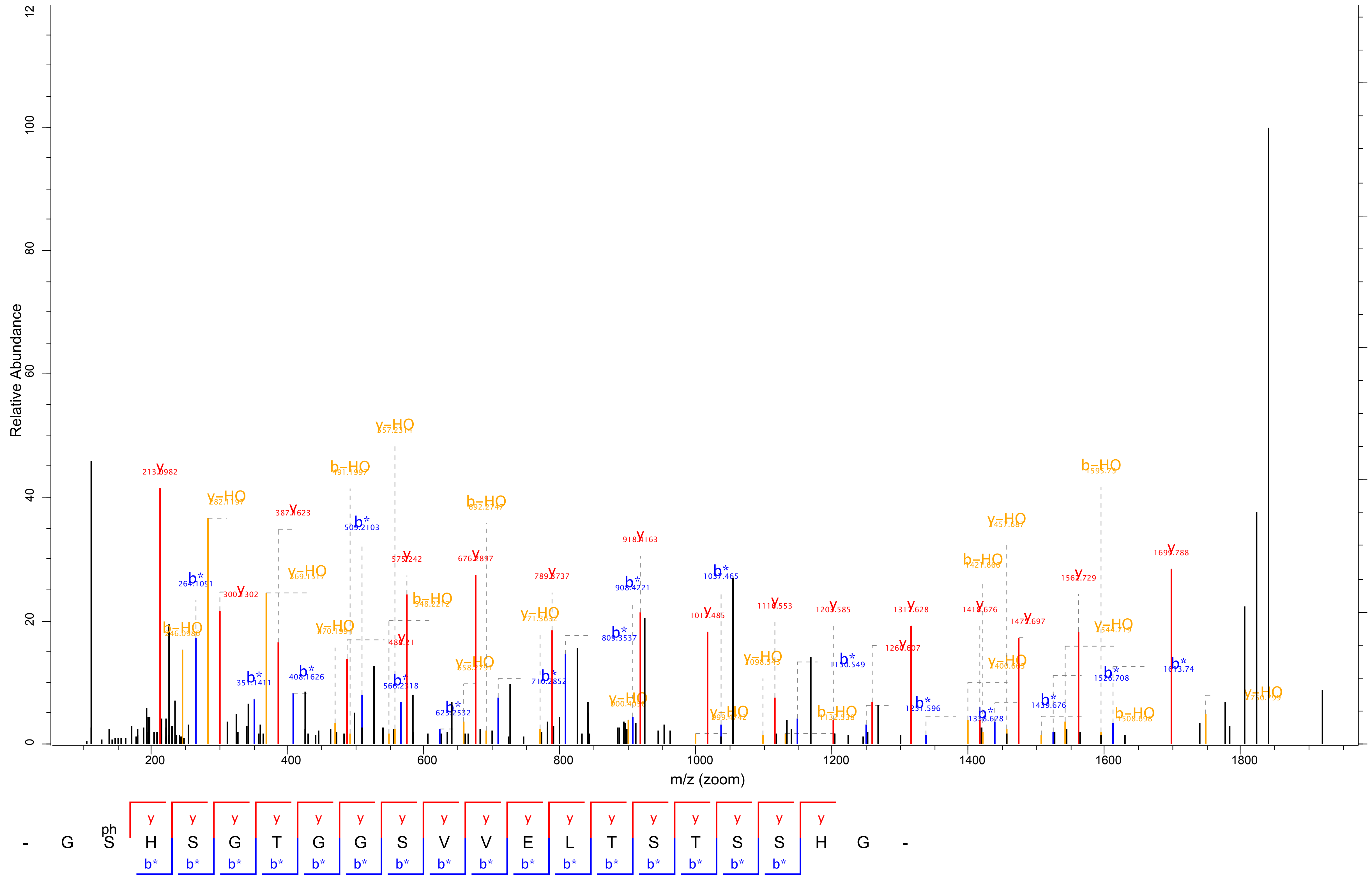

| Raw File | Scan | Method | Score | m/z |
| --- | --- | --- | --- | --- |
| P3_2_2 | 7494 | FTMS; HCD | 44.54 | 641.94 |

### pS827 by AGC2-2

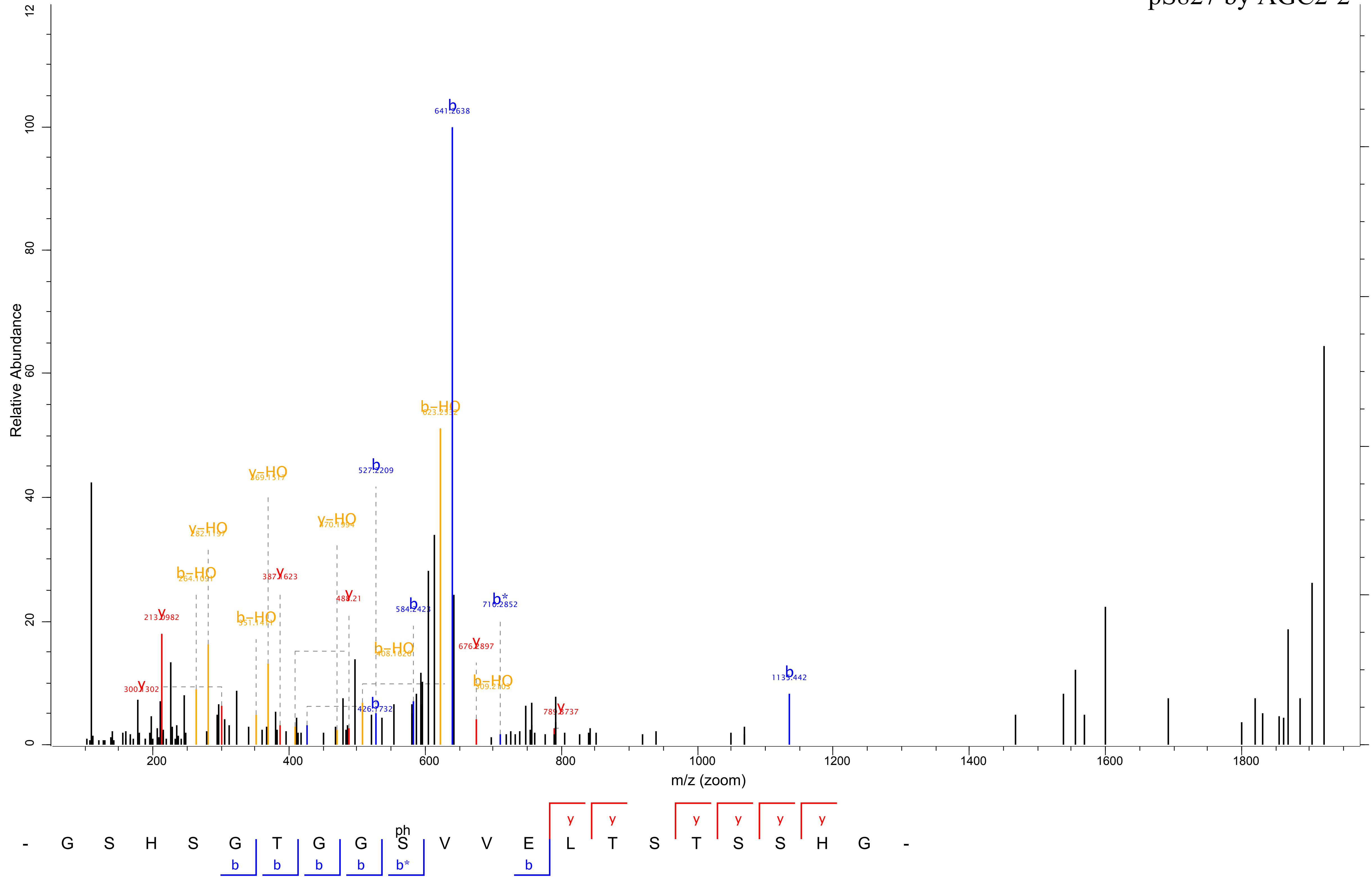
